# A common tolerance mechanism of bacterial biofilms to antibiotics

**DOI:** 10.1101/2023.01.30.526163

**Authors:** Yuzhen Zhang, Yumin Cai, Tianmin Wang, Qile Wu

## Abstract

A deeper mechanistic understanding of antibiotics tolerance exhibited by bacterial biofilms is crucial for clinical treatment and drug development. Here, we found a common binary distribution for various antibiotics in biofilms, which was determined by glucose distribution. Antibiotic penetration would be accompanied by glucose leakage. Surprisingly, the biofilm periphery exhibited constant glucose consumption and antibiotics patterns after drug treatment, while planktonic bacteria did not. Spatial multi-omics revealed that activation of lipid metabolism was responsible for continuous glucose consumption, which was channeled to thicken the cell membrane and subsequently weakened antibiotic penetration, leading to a low drug concentration in the biofilm interior - a common mechanism underlying biofilm drug tolerance. We further revealed that the biofilm-specific activation of lipid metabolism was derived from slower drug penetration and longer response time of biofilms to antibiotics, owing to lower energy metabolism and membrane potential in biofilms compared to planktonic bacteria.

## INTRODUCTION

Bacteria that form aggregates and develop as biofilms exhibited significantly enhanced tolerance against killing by antibiotics compared with the planktonic bacteria ^1–3^. Reduced antibiotic susceptibility contributes to the persistence of biofilm infections such as those associated with implanted devices, which causes great challenges to the eradication of such infections ^1, 4, 5^. The exceptional tolerance of biofilm against antibiotic treatment works largely at the phenotypic level. For example, bacteria that lack protective mutations or that lack other mobile genetic elements carrying resistance genes, nevertheless become less susceptible when grown in the biofilm state ^6, 7^. Antibiotic sensitivity is usually quickly restored when bacteria are dispersed from a biofilm ^6^; Such fast and reversible switch of tolerance level between biofilm and dispersed planktonic bacteria therefore suggests an adaptive tolerance mechanism at phenotypic level rather than a genetic alteration at this stage ^8^. Moreover, phenotypic tolerance at the first stage gives rise to a high number of bacterial cells that survive antibiotic treatment, which is essential to the potential evolution of genetic resistance ^9^; And such evolution process could be further accelerated or diversified by the drug concentration gradient in biofilms ^10, 11^. Hence, understanding the molecular mechanism of the unique and extraordinary phenotypic drug tolerance in biofilm is critical to the combating against the prevalent biofilm-associated infectious diseases.

As an emergent phenotype in a bacterial community, biofilm drug tolerance at phenotypic level is assumed to depend on some unique characteristics of the community, such as poor antibiotic penetration ^12–15^, nutrient limitation and slow growth of subpopulations ^12, 16–19^. Basically, these previous researches regard biofilm drug tolerance from a static point of view — some unique characteristics in biofilm exist in prior to drug treatment and are responsible for the enhanced tolerance, including extracellular matrix (ECM) that inhibits drug diffusion ^12–15^, spatial heterogeneity that results in drug-unsusceptible dormant subpopulations, etc.^5^ Despite the fact that biofilm drug tolerance could be partially attributed to these static features, extraordinary tolerance still existed in the absence of these assumed community characteristics. For example, fluoroquinolones, rifampicin, and ampicillin can penetrate the ECM well, but still cannot effectively eradicate the biofilms ^6, 20, 21^. Additionally, our previous work showed that the nutrient-limited interior region in the biofilm was not that metabolically dormant as previously assumed, due to the enhanced cross-regional metabolic coordination (Nature Chemical Biology, In press). In this context, we hypothesized that some community-unique dynamic features beyond those previously addressed static ones may functionally contribute to the emergent drug tolerance in biofilms. In particular, biofilm is a spatially structured community with heterogenous subpopulations in space ^12, 16–19, 22^. Dynamic responses of these subpopulations to drugs and their interactions in this process are poorly explored due to technical limitations; And these dynamic features may act as another layer of defense commonly against a wide range of antibiotics.

To understand the potential contribution of dynamic drug response to antibiotic tolerance in biofilm, a key gap is the lack of quantitative monitoring of real-time community characteristics, including antibiotics enrichment and adaptive gene expression with spatial resolution. Here, we combined several new techniques developed in our lab, including quantitative microfluidics for biofilm culturing, time-lapse microscopy to monitor drug response and spatial multi-omics methods for biofilms, to dissect drug distribution and biofilm response in space and time. Using these methods, we uncovered a dynamic mechanism deployed by various biofilms to protect the interior region from the killing of many different antibiotics. This mechanism depends on first, the poor drug efficacy against the periphery region due to the inherently depolarized membrane in biofilm; and second, the resulting time window for the biofilm periphery to maintain constant glucose consumption and adopt essential adaptations; third, the activated lipid metabolism to thicken bacterial membrane and further limit drug diffusion. This dynamic process give rise to a universal protection of the interior region in biofilms; And our work also uncovers a new paradigm that drug tolerance in biofilm emerges from the dynamic interactions between physiologically divergent subpopulations within a spatial scale of only hundreds of micrometers.

## RESULTS

### Glucose gradient determines the binary distribution of antibiotics in biofilms

Characterizing the intracellular concentrations of antibiotics is a key step in quantifying their efficacy. Among the common antibiotics, a few antibiotics, such as tetracycline (TET), berberine (Berb), and chromomycin A3 (Chromo A3), are intrinsically fluorescent; thus, it is possible to use auto-fluorescence to characterize their intracellular concentration by using fluorescence microscopy in real time and without incurring damage. However, most antibiotics (such as kanamycin (KAN), ciprofloxacin (CIP), and ceftazidime (CEF)) do not exhibit fluorescence or exhibit weak fluorescence; therefore, it is challenging to characterize their effective intracellular concentration in real-time and situ. Fortunately, Kohanski et al. ^23^ reported that three major classes of bactericidal antibiotics (aminoglycosides, β-lactams, and quinolones) caused an increase in intracellular reactive oxygen species (ROS) when these antibiotics enter the cell, and the ROS level can be measured using dichloro-dihydro-fluorescein diacetate (DCFH-DA) fluorescent dye ^24^. Therefore, for these antibiotics, the fluorescence intensity of DCFH-DA may be used to characterize their intracellular concentrations.

Based on microfluidics combined with time-lapse microscopy for quantitatively studying biofilms previously developed by our team ^25^, we found that fluorescent antibiotics (TET, Berb, and Chromo A3) presented heterogeneous fluorescence distribution in *Escherichia coli* biofilms (Figure 1A). TET fluorescence was more concentrated at the periphery of the biofilm than in the interior, with a sharp decrease in fluorescence visible between the periphery and the interior. Berb and Chromo A3 presented patterns opposite to TET, displaying a higher intensity in the interior than that in the periphery. Further quantitative characterization by high-performance liquid chromatography (HPLC) showed that the intracellular concentration of TET in the biofilm periphery was approximately twice that in the interior (Figure 1B; refer to the STAR Methods regarding the segmentation of the biofilm into the periphery and interior), while the intracellular concentration of Berb in the biofilm interior was three times that in the biofilm periphery. Both results agreed quantitatively with those of the fluorescence ratios (Figure 1C; refer to Figure S1 and STAR Methods for the calculation of spatial profile and temporal profiles), indicating that the fluorescence intensity of such antibiotics reflects their intracellular drug concentration, and the heterogeneous distribution of fluorescence represents the heterogeneous action of these antibiotics in biofilms.

**Figure 1:**
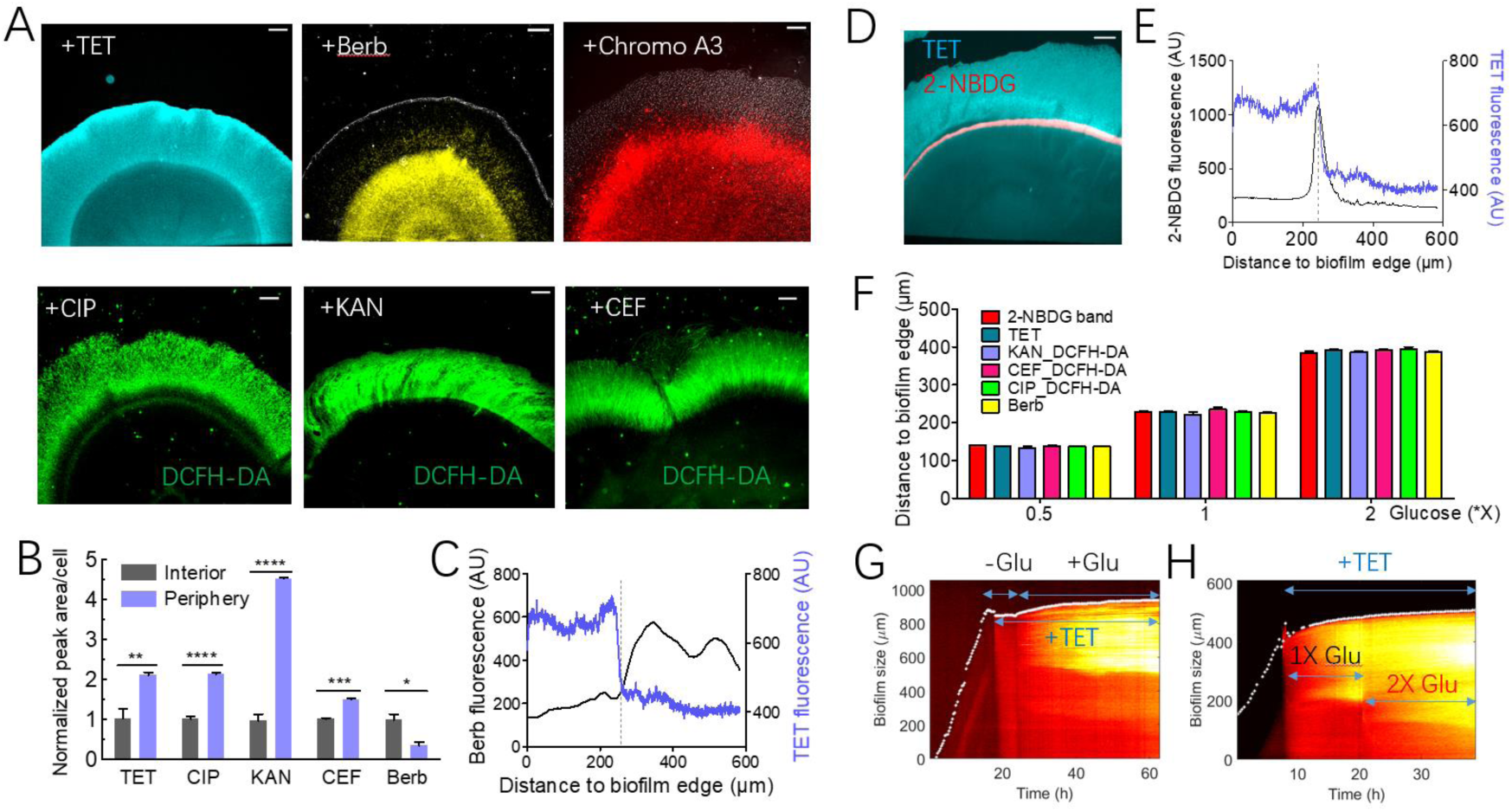
Spatial distribution of antibiotics in biofilms is determined by glucose distribution. (A) Fluorescence distribution of antibiotics in biofilms. TET (TET), Berberine (Berb) and Chromomycin A3 (Chromo A3) are fluorescent antibiotics. Kanamycin (KAN), ceftazidime (CEF), and ciprofloxacin (CIP) are bactericidal antibiotics; they all produce hydroxyl radicals which make the DCFH-DA dye fluoresce. Antibiotics concentrations were as follows: 5 μg/ml TET, 200 μg/ml Berb, 20 μg/ml Chromo A3, 0.06 μg/ml CIP, 10 μg/ml KAN, and 4 μg/ml CEF. TET, CIP, KAN, and CEF caused growth arrest after addition, thus snapshots showed the scene at 20 h after them addition. Berb and Chromo A3 had no impact on biofilm growth, thus they were added in the medium initially. Scale bar, 100 μm. (B) Intracellular antibiotics concentration in biofilms quantified by HPLC and LC-MS. *n*=3. Statistical significance was determined by two-sided Student’s t-tests: *, *p* < 0.05; **, *p* < 0.01; ***, *p* < 0.001, ****, *p* < 0.0001. (C) Spatial distribution of TET is spatially complementary to that of Berb. (D and E) The TET band spatially coincided with the 2-NBDG band. (F) Antibiotics distribution changed with glucose concentration. (G and H) TET distribution changed with glucose distribution.

The fluorescence of DCFH-DA becomes enhanced after it reacts with intracellular ROS, and antibiotics such as KAN, CIP and CEF caused an increase in intracellular ROS levels ^23^. Hence, using DCFH-DA signal as an indicator, we found that KAN, CIP, and CEF, showed significantly higher DCFH-DA fluorescence in the biofilm periphery than in the interior (Figure 1A), suggesting that these antibiotics mainly enriched in and act on the biofilm periphery. Indeed, through quantitative characterization by HPLC (used for CEF and CIP) and liquid chromatography–mass spectrometry (LC-MS; used for KAN), we found that the intracellular concentration of KAN, CIP and CEF in biofilm periphery was about 1.8-4.5 times that in interior (Figure 1B). Therefore, the DCFH-DA fluorescence pattern can be reliably used to characterize the spatial patterns of antibiotics that cause ROS production in real-time and situ.

Based on the characterizations above, we found that antibiotic distribution in biofilms could be summarized into two localization patterns: TET, KAN, CIP, and CEF were distributed in the biofilm periphery, whereas Berb and Chromo A3 were distributed in the biofilm interior (Figure 1A). Notably, these two patterns appeared highly complementary in space: TET and related antibiotics were distributed in the 220 µm periphery of the biofilm, while the distribution of Berb and Chromo A3 were exactly enclosed by this periphery region (Figures 1A and 1C). Such complementary mode suggested that it was one intrinsic metabolic characteristic of the biofilm that determined these spatial patterns of antibiotic distribution. Agreeing with this hypothesis, we found that the spatial distribution of antibiotics was consistent with that of carbon sources (Figures 1D-1F). Briefly, as glucose was the only carbon source in the medium, we used the glucose analog, 2-NBDG, as a fluorescence marker of glucose depletion ^23, 25^ (STAR methods); Accordingly, the region within the fluorescence 2-NBDG band indicates glucose starvation (Figure 1D). Indeed, the TET boundary (between the high- and low-signal region) coincided with the 2-NBDG band (Figures 1D and 1E). Furthermore, the distribution of various other antibiotics also depended on glucose distribution (Figure 1F); Altering the glucose concentration could alter the glucose penetration depth, as well as the spatial pattern of antibiotics (Figure 1F). Using TET as an example, we demonstrated that TET failed to infiltrate the biofilm and exhibited no spatial enrichment when glucose was absent in the medium. Upon restoring 1x (22 mM) glucose, TET rapidly and synchronously penetrated throughout the glucose band (Figures 1G and S2A). Switching from 1x to 2x glucose exhibited similar results (Figures 1H and S2B). These results shows that glucose distribution and derived energy metabolism gave rise to the spatial patterns of antibiotic enrichment.

### Biofilm periphery maintains glucose consumption under antibiotic stress, whereas planktonic cells do not

For most of the antibiotics (KAN, CIP, CEF, TET), higher enrichment at the periphery region was observed. This is in sharp contrast to those enriched at the interior region, such as Berb and Chromo A3 (Figure 1A), which could be simply attributed to poor efflux activity there due to glucose starvation and inhibited energy metabolism ^25^. The enrichment of antibiotics at the periphery region is functionally profound to biofilm, as it provides protection towards the interior region upon antibiotic stress. Indeed, we observed unchanged translational activity at the interior region of the biofilm under TET treatment (Figures S2C and S2D); And confocal microscopy suggested that TET fluorescence was mainly intracellular in the biofilm periphery, and extracellular in the biofilm interior (Figure S2E). Moreover, we excluded the contribution of ECM barrier to such protection, as the change of TET fluorescence was spatially synchronous in biofilm after glucose removal (Figures S2F and S2G). Therefore, we next sought to explore the mechanism underlying such protection uniquely for the interior region in biofilm when challenged by many clinically relevant antibiotics.

Given that glucose distribution or accessibility determined the distribution of antibiotics (TET, KAN, CIP, and CEF) (Figure 1), antibiotics would penetrate into the biofilm interior once glucose penetration occurred. In such context, the protection towards the interior region would be disrupted. However, we actually observed quite stable spatial patterns for those antibiotics enriched at periphery region for days (approximately 30 h, Figures S2H-2K). To resolve this contradiction, we measured the energy metabolism in space using 2-NBDG indicator after antibiotic treatment. Upon the addition of TET, KAN, CIP, CEF, rifampicin (RIF), or trimethoprim (TMP) (2x the minimal inhibitory concentration (MIC)), biofilm growth quickly stopped (Figure 2A); but to our surprise, the boundary of glucose starvation in biofilms did not change significantly after the addition of antibiotics, as indicated by the 2-NBDG band (Figures 2A and 2B). This result was further confirmed by measuring glucose concentration in the microfluidic waste (Figure 2C). In sharp contrast to this result, we indeed observed completely halted glucose consumption in planktonic bacteria (Figure 2D). Taken together, these results demonstrated that the biofilms could maintain peripheral glucose consumption, impede glucose and antibiotics penetration into the interior, and exhibit a community-specific mechanism for maintaining antibiotic tolerance.

**Figure 2:**
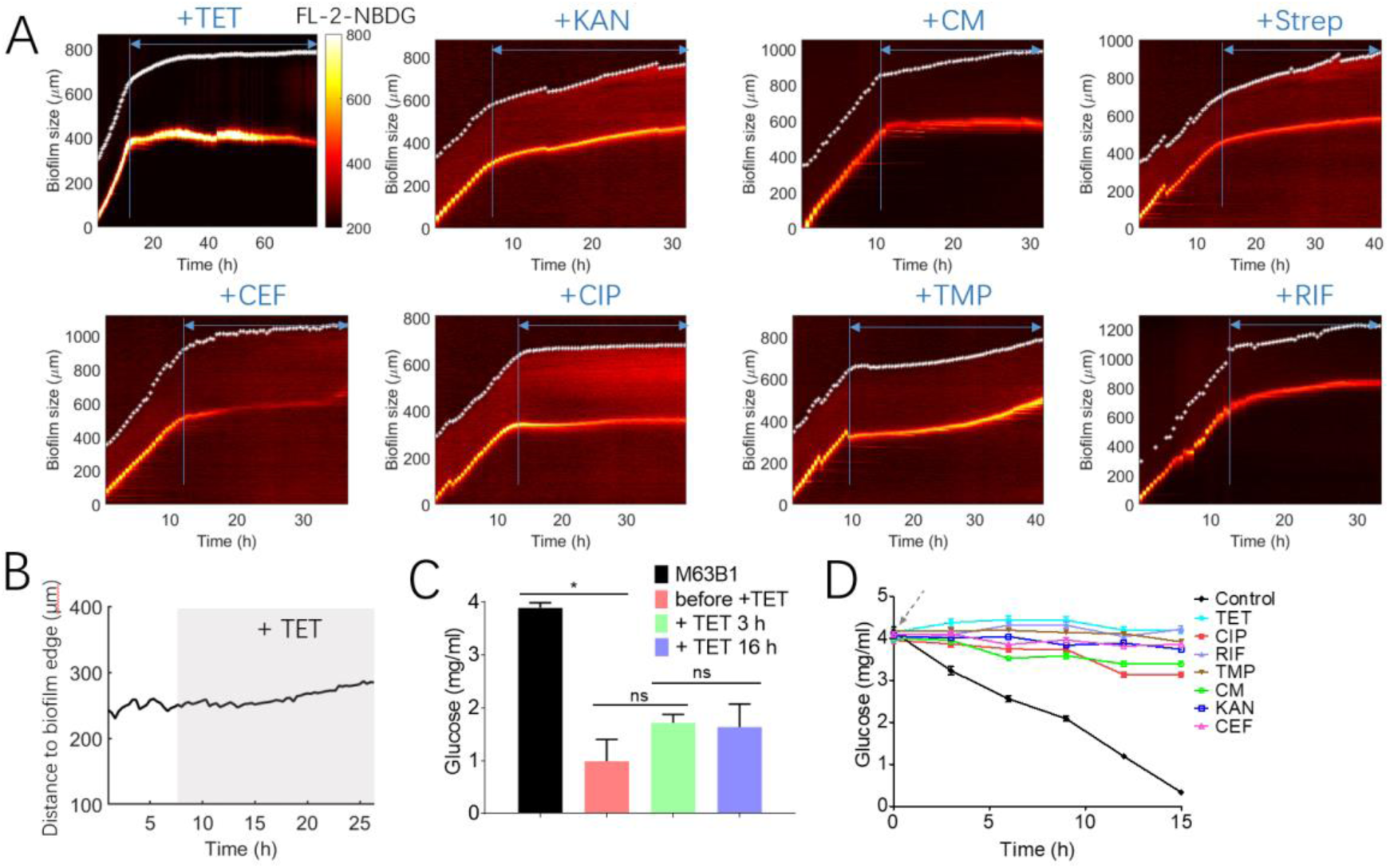
Biofilm periphery maintains glucose consumption under antibiotics stress, but planktonic cells do not. (A) 2-NBDG band before and after the addition of various antibiotics to biofilms. Antibiotics concentrations were as follows: 5 μg/ml TET (TET), 10 μg/ml kanamycin (KAN), 14 μg/ml chloramphenicol (CM), 10 μg/ml streptomycin (Strep), 4 μg/ml ceftazidime (CEF), 0.06 μg/ml ciprofloxacin (CIP), 3 μg/ml trimethoprim (TMP), and 20 μg/ml rifampicin (RIF). (B) 2-NBDG band was almost unchanged after TET addition. (C) Glucose concentration in the waste of the chip before (pink column) and after TET addition (green and purple columns). The black column presents the inlet concentration of glucose in the chip. *n* = 3. Statistical significance was determined by two-sided Student’s t-test:s *, *p* < 0.05; **, *p* < 0.01; ***, *p* < 0.001; ns, not significant. (D) Glucose concentration in the supernatant of planktonic bacteria culture after the addition of antibiotics. Zero time represents when antibiotics were added. The concentrations of antibiotics added were equal to those in biofilms.

### Spatial multi-omics reveal unique activation of lipid metabolism responsible for glucose consumption in biofilm periphery under antibiotics stress

Given that biofilm growth was stopped but glucose consumption was maintained upon the abovementioned antibiotic treatment, we wondered where the generated energy was diverted to in metabolic network, which probably contributed to the unique tolerance to these antibiotics in biofilm. To answer this question, we developed a spatial multi-omics method for biofilms (Figure 3A, STAR Methods), via upgrading our previous design ^25^ to a new omics chip (L×W×H = 30 mm×6 mm×100 μm) suitable for omics sample preparation. The number of bacterial cells in the biofilm reached 100 million, and the total protein amount extracted from a single biofilm was approximately 50 μg. Cell number and protein amount enabled sample preparation in metabolomics and proteomics. Using the fluorescence indication of the 2-NBDG band, we quickly separated the peripheral and interior regions of a biofilm along the 2-NBDG band under a stereofluoroscope (Figure 3A, STAR Methods), which enabled profiling proteomes, intracellular metabolomes or lipidomes from these two regions in biofilm with different physiological states.

**Figure 3:**
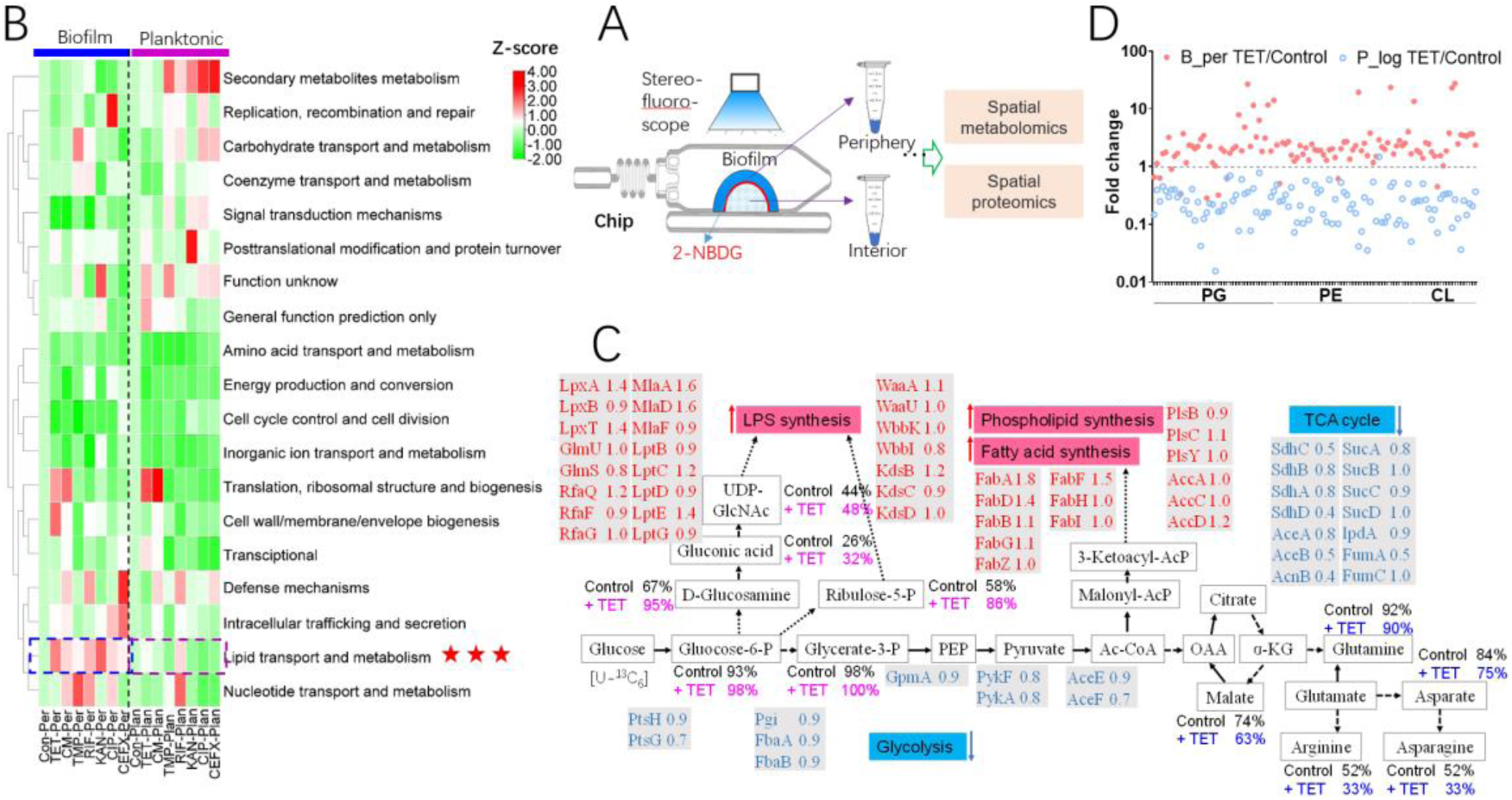
Spatial multi-omics reveal lipid metabolism is upregulated in biofilms under stress from various antibiotics. (A) Workflow of spatial proteomics and spatial metabolomics in biofilms. (B) Overview of the spatial proteomics between the *E. coli* biofilm and planktonic cell before and after the addition of various antibiotics. The fraction of total protein massed assigned to the 19 Clusters of Orthologous Genes (COG) categories is indicated for each condition. For the biofilm periphery, the COGs after the addition of antibiotics for 20 h are normalized with the control (before addition of antibiotics), and the same was done for planktonic bacteria. Then, all COGs were normalized by Z-score, for which a color bar is shown. Antibiotics concentrations were as follows: 5 μg/ml tetracycline (TET), 10 μg/ml kanamycin (KAN), 14 μg/ml chloramphenicol (CM), 4 μg/ml ceftazidime (CEF), 0.06 μg/ml ciprofloxacin (CIP), 3 μg/ml trimethoprim (TMP), and 20 μg/ml rifampicin (RIF). (C) Fold change of proteins (biofilm periphery +TET/control) in central carbon metabolism (in gray shadow). Allocation of carbon atom with (+TET) and without (Control) the addition of TET (5 μg/ml, 20 h). Glucose (U-^13^C_6_) replaced for 10 h. (D) Lipidomics change in *E. coli* biofilm and planktonic cells before and after the addition of TET. PE, phosphatidylethanolamine; PG, phosphatidylglycerol; CL, cardiolipin. UDP-GlcNAc, uridine diphosphate N-acetylglucosamine; PEP, phosphoenolpyruvic acid; Ac-CoA, acetyl coenzyme A; OAA, oxaloacetic acid; α-KG, α-ketoglutarate; LPS, lipopolysaccharide; TCA, tricarboxylic acid.

We used data-independent acquisition (DIA), label-free, quantitative proteomics to detect the proteome. We detected 2,600 proteins in *E. coli* BW25113 biofilm and planktonic bacteria (Data set 1 in Supplementary Table 1). This is slightly higher than the number of detected proteins (2,300) in a benchmark study ^26^, suggesting the sensitivity of our pipeline. We filtered 54 sets of data using a common threshold (STAR Methods), giving rise to 1,908 proteins detected in all these samples (Data set 2 in Supplementary Table 1). Based on the Clusters of Orthologous Genes (COGs) ^27, 28^ results (Data set 3 and 4 in Supplementary Table 1), we found that the lipid transport and metabolism pathway was uniquely activated in the periphery of the biofilm after treatments by all tested antibiotics (Figure 3B, TET, KAN, CIP, CEF, RIF, and TMP), regardless of the bactericide, bacteriostat, or different targets of action. Notably, lipid metabolism was almost completely inhibited in planktonic bacteria (Figure 3B), indicating a unique activation of lipid metabolism at biofilm periphery when challenged by antibiotics. A more detailed functional enrichment analysis (Figure S3, STAR Methods) suggested that after 20 h of TET treatment, parts or all of the phospholipids, LPS, and fatty acid synthesis pathways were activated. Actually, the viability of all gram-negative bacteria depends on the maintenance of a tight balance between phospholipids and LPS ^29^, and biosynthesis of both these components uses a common metabolic precursor, fatty acyl-ACP, a key intermediate molecule from the fatty acid biosynthesis pathway ^30^.

To further explore the impact of antibiotic treatment on the physiology of biofilm periphery region, especially its lipid metabolism, we integrated data from spatial proteomics and C13 metabolomics (Figure 3C) under TET treatment. From the results of metabolic flux, we found that glucose in the periphery of the TET-treated biofilm flowed more into the ribulose-5-P and UDP-glucosamine than it did in untreated cells (Figure 3C), both of which were critical precursors for the biosynthesis of sugar component in LPS. Consistently, proteomics revealed that glycolysis and tricarboxylic acid cycle (TCA) pathways were downregulated, while the pentose phosphate pathway (PPP) was enhanced to supply ribulose-5P. Moreover, fatty acid and the subsequent phospholipid biosynthesis were upregulated to supply both the lipid part of LPS and cell membrane (Figure 3C). In contrast, amino acids biosynthesis were significantly downregulated (Figures 3B and 3C). Further, through spatial lipidomics, lipids in planktonic cultures were downregulated following TET treatment, while the lipid content at the biofilm periphery increased significantly (Figure 3D). The above results indicate that antibiotic treatment significantly activated lipid synthesis at the biofilm periphery, which is primarily responsible for continuous glucose consumption.

### Activation of lipid metabolism thickens the cell membrane and weakens antibiotic penetration, leading to biofilm tolerance to antibiotics

Next, we wondered the relationship between activated lipid metabolism at the periphery region and drug tolerance in the biofilm. Using a highly lipophilic dye — FM 4-64, which stains membrane lipids ^31^, we found that the peripheral fluorescence of FM 4-64 was enhanced after TET treatment, exhibiting a pattern similar to that of TET (Figure 4A). Furthermore, using confocal microscopy (Figures 4B, 4C, and S4A), at the single cell level, we found that the thickness of the lipid envelope of peripheral bacteria in biofilm increased by approximately two-fold following TET treatment for 20 h, when compared to that without TET treatment (Figure 4C). Further, using transmission electron microscope (TEM), we revealed that the thickness of the bacterial outer membrane (Figure S4B) increased by approximately two-fold after 20 h and 60 h of TET treatment, when compared to that without TET treatment (Figures 4D, 4E, and S4B).

**Figure 4:**
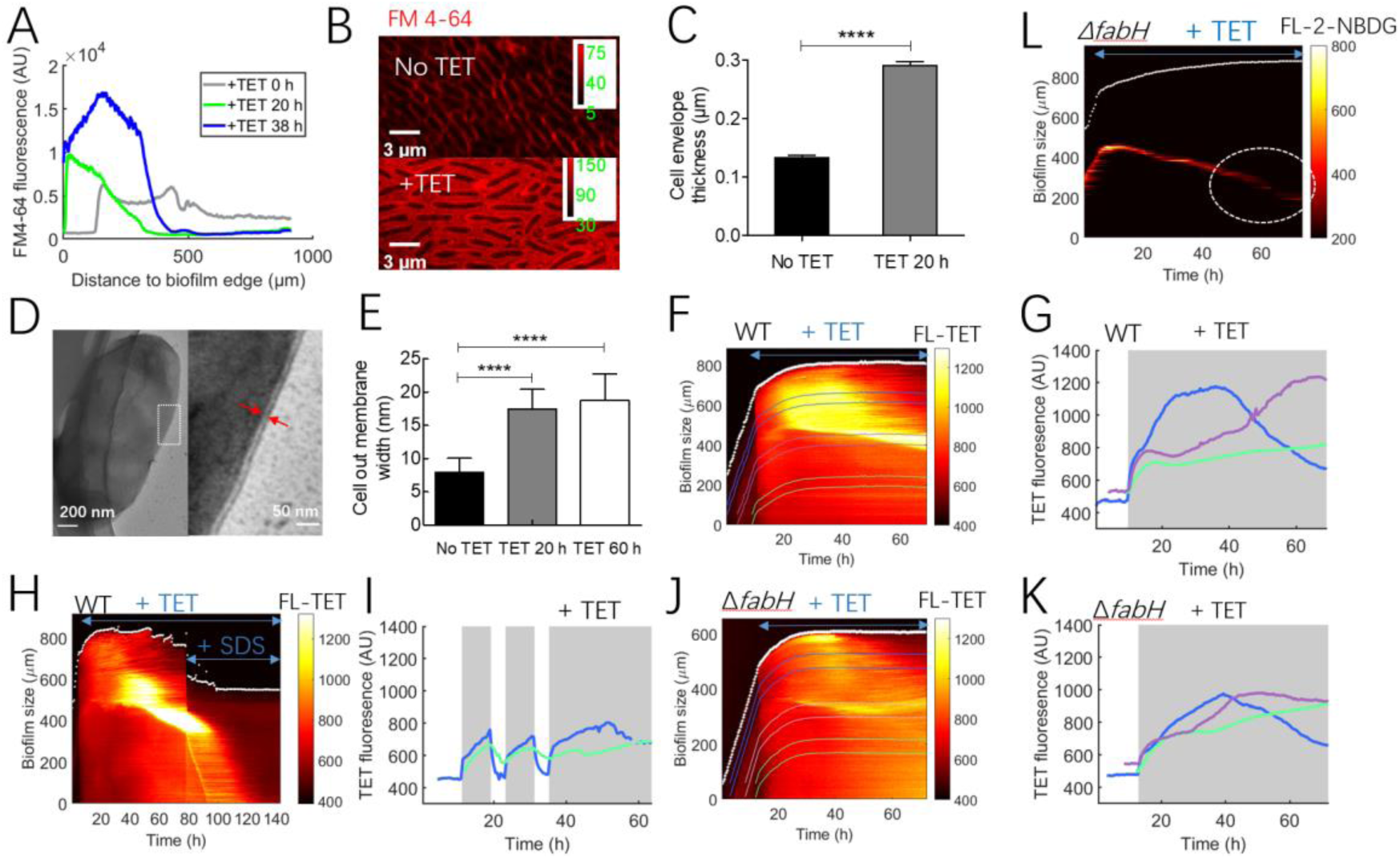
Activation of lipid metabolism thickens the cell membrane, thus weakening the penetration of antibiotics. (A) Spatial profile of FM 4-64 in the biofilm after the addition of tetracycline (TET) for 0, 20 and 38 h. (B) Confocal images of the cells in the biofilm periphery before and after addition of TET (20 h). (C) Cell envelope thickness before and after the addition of TET. *n* = 52. (D) Transmission electron microscopy images of the cells in the biofilm periphery. The red arrow indicates the bacterial outer membrane. (E) Outer membrane thickness before (n = 266) and after addition of TET (20 h (*n* = 119) and 60 h (*n* = 92)). (F) Kymograph of TET pattern in WT *E. coli* biofilm. Five μg/ml TET was added. (G) Temporal profile of TET accumulation in periphery (blue line), middle (magenta line) and interior (green line) of the biofilm in (F). (H) Kymograph of TET pattern in WT *E. coli* biofilm after addition of 0.03% sodium dodecyl sulfate (SDS). (I) TET fluorescence accumulation in biofilm (periphery, blue line; interior, green line) when TET was added periodically shown in Figure S5B. (J) Kymograph of TET pattern in Δ*fabH E. coli* biofilm. (K) TET fluorescence accumulation in the Δ*fabH* biofilm shown in (J). (L) 2-NBDG band before and after the addition of TET in Δ*fabH* biofilm. Statistical significance in relevant panels was determined by two-sided Student’s t-tests: *, *p* < 0.05; **, *p* < 0.01; ***, *p* < 0.001, ****, *p* < 0.0001.

Studies have shown that increasing the length of the phospholipid acyl tails thickens the hydrophobic layer of the membrane ^32^. The permeability coefficient of a molecule is inversely proportional to the membrane thickness. For tail lengths from 14 to 26 carbon atoms, elongating the lipid tails by two carbon atoms decreases the permeability coefficients by a factor of approximately 1.5 ^32^. Moreover, it is previously found that bacteria modify their membrane composition in response to environmental changes, such as in temperature, osmolarity, salinity, and pH, giving rise to enhanced tolerance ^33–35^. In particular, many pathogens alter their membrane phospholipid structures to increase their resistance to antimicrobial peptides ^36^; High temperatures and envelope stress also increased LPS synthesis ^30^. Given that the lipid content was significantly increased (Figure 3D) and the length and saturation of the C-chains were both enhanced (Supplementary Table 2), we hypothesized that the lipid metabolism activation and cell membrane thickening would weaken drug penetration at the biofilm periphery. Following TET treatment, a bright periphery and dark interior pattern were maintained for approximately 40 h in wild-type (WT) biofilms (Figure 4F). After 60 h, even if the peripheral bacteria died, a bright TET band appeared in the middle, with a low fluorescence intensity still robustly maintained in the interior (Figures 4F and 4G); We inferred that this band replaced the barrier function of periphery region to block drug penetration into the biofilm interior at this stage. Indeed, the average thickness of the outer membrane slightly increased at 60 h of TET treatment compared with that at 20 h (Figures 4E and S4B), characterized by TEM. When a membrane detergent (sodium dodecyl sulfate (SDS) ^37^, no impact on biofilm growth (Figure S5A)) was used to destroy this assumed protective layer, the TET quickly entered the biofilm interior (Figure 4H). Moreover, if we add SDS prior to TET, the TET pattern was just maintained for less than 15 h, then TET penetrated into the interior of the biofilm (Figure S5A). Based on this finding, we predicted that the TET penetration rate would be further reduced if lipid expression was induced in advance. Indeed, after 8 h of TET pre-exposure, the penetration rate of TET in the peripheral decreased, and TET concentration in the interior was almost flattened and no longer increased after further prolonged TET treatment (Figures 4I and S5B). Other pre-exposure at different times caused similar results (Figures S5C-5F).

We next sought to block lipid synthesis activation by genetic manipulation to validate our hypothesis. In particular, via *fabH* knockout (the first step for fatty acid elongation, catalyzed by FabH ^36^, which play a role in governing the total rate of fatty acid ^38^ and membrane phospholipids production ^39^), the biofilm periphery could no longer continuously block TET penetration, resulting in a reversed TET pattern in space: the TET accumulation at interior region exceeded that of the periphery region, and the pattern disappeared after 40 h of TET treatment (Figures 4J and 4K). Consistently, peripheral bacteria could not maintain glucose consumption anymore after 40 h of TET treatment in the *ΔfabH* biofilm, as indicated by the disappearance of 2-NBDG band (Figure 4L). In contrast, 2-NBDG band indicated that glucose consumption persisted in WT biofilm for up to 65 h or even longer after TET treatment (Figure 2A).

Akin to TET, various antibiotics universally induce lipid metabolism (Figure 3B); therefore, we assumed that activated lipid metabolism led to biofilm tolerance towards a wide range of antibiotics. Indeed, we found that after blocking lipid synthesis via *fabH* knockout, the biofilm tolerance to CIP, CEF, streptomycin (Strep), and TMP was all reduced, indicated by the enhanced efficacy of these antibiotics upon the interior region monitored by ROS production (Figures 5A and 5B). This finding indicates that the activation of lipid metabolism plays an important role in protecting the interior biofilm from the killing by these antibiotics. In addition, we found the localization patterns of TET in *Klebsiella pneumoniae* and *Salmonella typhimurium* biofilms were similar to those observed in *E. coli*, with a bright periphery and a dark interior, maintaining this pattern for 40 h or longer (Figure 5C). Akin to *E. coli*, following *ΔfabF* knockout in *K. pneumoniae* and *ΔfabH* in *S. typhimurium*, which both interrupted fatty acids biosynthesis, the patterns of bright periphery and dark interior were both partially challenged---TET was easier to penetrate into biofilms formed by knockout strains (Figure 5C). Taken together, these results revealed that the maintained glucose consumption supported the activation of lipid metabolism at the periphery region of the biofilm; This mechanism not only protects the periphery itself, but also prevents more fatal antibiotic penetration into the biofilm interior, which is a common tolerance mechanism deployed by various biofilms against a wide range of antibiotics.

**Figure 5:**
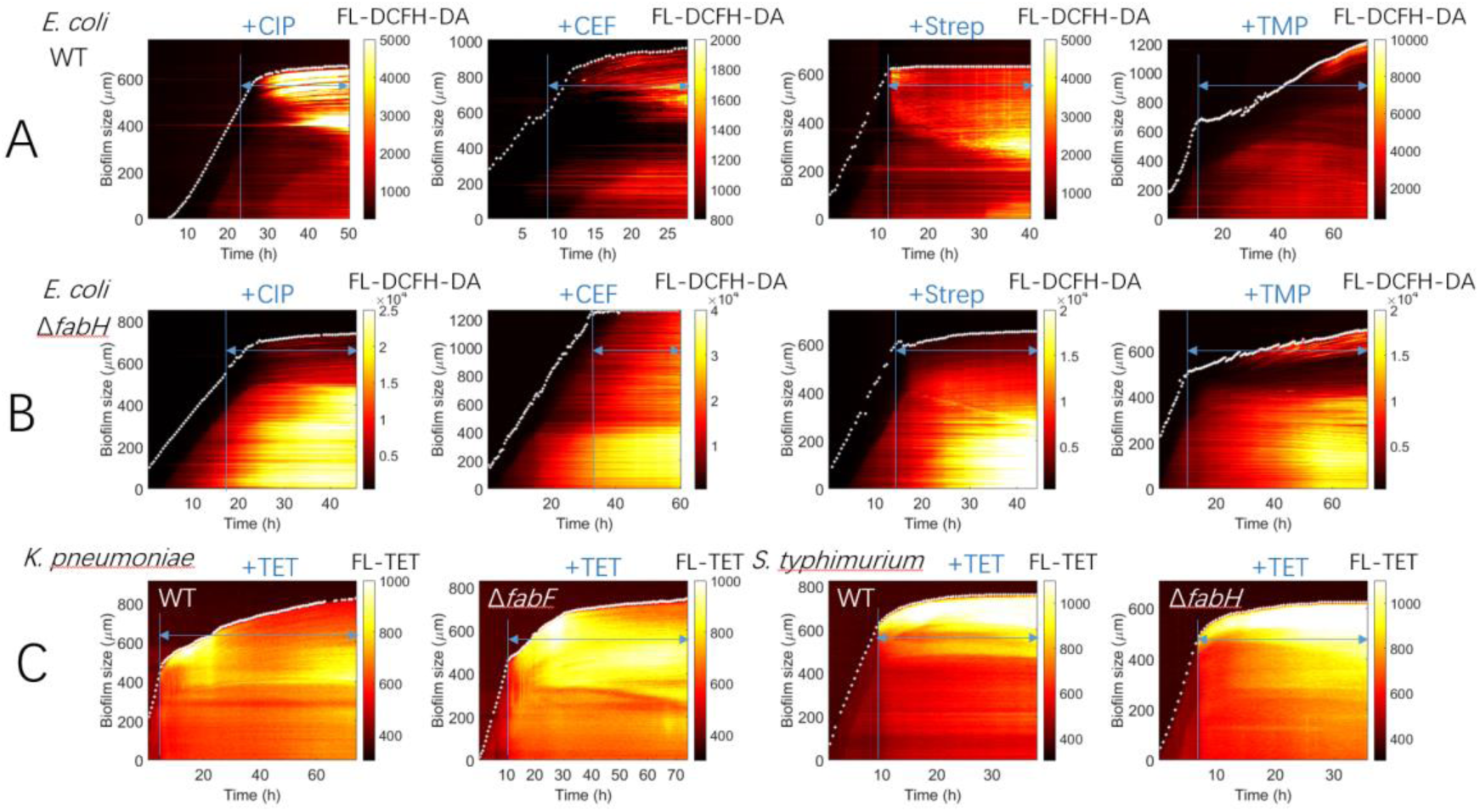
Lipid metabolism in different types of biofilms in response to various antibiotics. (A and B) Kymograph of DCFH-DA fluorescence accumulation in WT (A) and Δ*fabH* (B) *E. coli* biofilms before and after the addition of various antibiotics. 0.06 μg/ml ciprofloxacin (CIP), 3 μg/ml trimethoprim (TMP), 4 μg/ml ceftazidime (CEF), and 10 μg/ml streptomycin (Strep) were added. (C) Kymograph of tetracycline (TET) pattern in *K.pneumoniae* (WT and Δ*fabF*) and *S. typhimurium* (WT and Δ*fabH*) biofilms. 6 μg/ml TET was added.

### Low membrane potential in biofilm reduces the initial penetration of antibiotics and creates a time window to respond to antibiotics

A fundamental question remains unanswered: why do biofilms, but not planktonic bacteria, maintain glucose consumption and activate lipid metabolism? The first evidence came from the accumulation dynamics of TET in biofilm and planktonic bacteria. In particular, we found that the time required for 5 μg/mL TET to accumulate to saturation at the periphery of the biofilm was approximately 18 h (Figures 6A and S6A). In contrast, the corresponding duration for planktonic bacteria was approximately 2 h, as determined by fluorescence-activated cell sorting (FACS) (Figure 6B). To further validate this result, we constructed an intracellular TET sensor, P*tetR*-*mCherry*, which initiates the expression of *mCherry* only when TET enters the cell and derepresses TetR. Following the addition of 1 µg/mL TET, the fluorescence of mCherry in the periphery of biofilm continued to increase for 18 h (Figure 6C), whereas the fluorescence of mCherry in planktonic bacteria in the same microfluidic chamber with biofilm reached equilibrium within 2 h (Figure 6D). TET is well known to inhibit protein translation; To exclude any effect caused by translation inhibition, we also characterized the penetration dynamics of 1 μg/mL anhydrotetracycline (aTC, an analogue of TET with no inhibition upon translation^40^), and the results were consistent with those of TET (Figures S6B and S6C). Therefore, TET accumulation in the biofilm was significantly slower than in planktonic bacteria. We reasoned that this difference could provide a time window for the biofilm to adopt its latent potential fighting against antibiotics, such as the activation of lipid metabolism; Functionally, such adaptation would inhibit drug penetration and gave rise to poorer drug efficacy. In line with this assumption, we showed that 5 µg/mL TET led to little (if any) membrane damage (determined by DiBAC4 (5) ^24, 41^, STAR Methods) when acting on biofilm (Figure 6E), but caused rapid and strong membrane damage to planktonic bacteria (Figure 6F).

**Figure 6:**
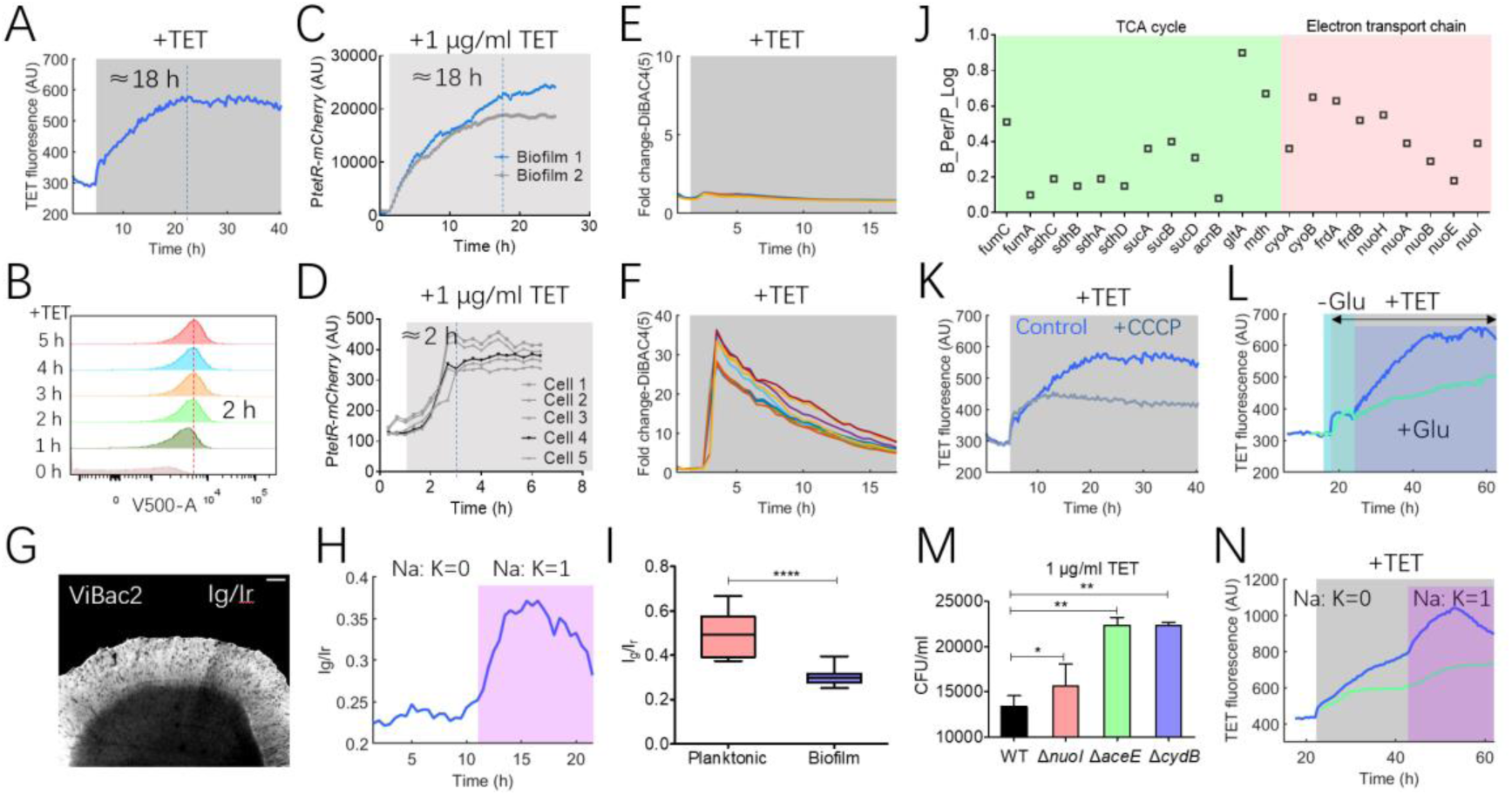
Low membrane potential in biofilms reduce the initial penetration of tetracycline (TET), creating a time window to respond to antibiotics. (A) Fluorescence of 5 μg/ml TET accumulation in *E. coli* biofilm shown in Figure S6A. (B) TET (5 μg/ml) accumulation rate in planktonic bacteria, as determined by fluorescence-activated cell sorting. (C and D) mCherry fluorescence in P*tetR*-mCherry biofilm (C) and planktonic cells (D) after the addition of 1 μg/ml TET, respectively. (E and F) DiBAC4(5) fluorescence in biofilm (E) and planktonic cells (F) before and after the addition of 5 μg/ml TET. (G) Membrane potential (Ig/Ir) distribution in *E. coli* biofilm. (H) Membrane potential (Ig/Ir) of cells in the biofilm increased significantly after changing the Na^+^: K^+^ ratio from zero to 1 shown in Figure S6E. (I) Membrane potential (Ig/Ir) in planktonic (*n* = 15) and biofilm bacteria (*n* = 24). Statistical significance was determined by two-sided Student’s t-tests: ****, *p* < 0.0001. (J) All detected proteins involved in energy metabolism in biofilm and planktonic bacteria. (K) Fluorescence of 5 μg/ml TET accumulation in *E. coli* biofilm periphery with or without 20 μM carbonyl cyanide m-chlorophenyl hydrazone (CCCP). (L) Fluorescence of 5 μg/ml TET accumulation in *E. coli* biofilm when glucose was removed and restored (periphery, blue line; interior, green line). (M) CFU/ml of WT, Δ*nuoI*, Δ*aceE and* Δ*cydB* cells (counted at 18 h) on a M63B1 plate with 1 μg/ml TET. *n* = 3. Statistical significance in relevant panels was determined by two-sided Student’s t-tests: *, *p* < 0.05; **, *p* < 0.01; ***, *p* < 0.001. (N) TET fluorescence increased significantly after changing the Na^+^: K^+^ ratio from zero to 1 after adding 5 μg/ml TET shown in Figure S7B (periphery, blue line; interior, green line).

To uncover the molecular mechanism underlying the difference of drug accumulation dynamics between biofilm and planktonic bacteria, we turned to the factors that may perturb drug uptake. It was reported that drug uptake such as aminoglycoside depended on the membrane potential ^42, 43^, a cellular characteristic with close relation with energy metabolism; And we found in this work that energy metabolism determined spatial drug accumulation in biofilm (Figure 1). Therefore, we wondered whether there existed inherent difference in membrane potential between biofilm and planktonic bacteria, to which the different drug accumulation dynamics could be attributed. To quantify spatial membrane potential pattern in biofilm, we adopted a recently reported membrane potential sensor ViBac2 ^44^. Firstly, we showed that the membrane potential at the periphery of the biofilm was higher than that the interior (Figures 6G and S6D), and this pattern was also consistent with the spatial pattern of energy metabolism (Figure S6D). Secondly, as reported in planktonic bacteria ^44^, switching the Na^+^: K^+^ ratio in the medium also caused cell membrane hyperpolarization in biofilm (Figures 6H, S6E-6G); And carbonyl cyanide m-chlorophenylhydrazone (CCCP; no impact on biofilm growth, see Figure S6H), an uncoupler of the proton gradient established during oxidative phosphorylation, could reduce membrane potential in the biofilm periphery (Figure S6I). These experiments suggested that this sensor could reliably probe the membrane potential in space within biofilm. Next, we adopted ViBac2 to quantify the membrane potential in biofilm and planktonic bacteria; Indeed, the membrane potential in planktonic single cells was approximately 60% higher than that in the biofilm periphery (Figure 6I), which may lead to more potent drug uptake and was thus consistent with our hypothesis. Interestingly, our spatial proteome data suggested that energy metabolism at the periphery region of biofilm was significantly lower than that in planktonic bacteria, in terms of proteins quantity in the tricarboxylic acid cycle (TCA) and the electronic transfer chain (ETC) (Figure 6J, Data set 4 in Supplementary Table 1); This result suggested that biofilm may inherently adopt some genetic programs to remodel its membrane potential, possibly in order to trigger higher tolerance towards a wide range of antibiotics.

After showing the difference in membrane potential between biofilm and planktonic bacteria, we next sought to validate its functional contribution to drug tolerance. Firstly, we showed that the pattern of higher periphery and lower interior of membrane potential in the biofilm (Figures 6G, S6D-6F) was highly consistent with the spatial pattern of energy metabolism (Figure S6D) and TET (Figure S6J). Second, CCCP could completely eliminate TET pattern in space (Figures 6K, S6K, and S6L). Third, if glucose was removed to decrease membrane potential in prior, the accumulation of TET in both periphery and interior quickly reached saturation with much lower absolute level (Figure 6L); Glucose addition then restored the TET accumulation dynamics at the periphery region until the normal state of saturation was reached (Figure 6L). Fourth, we adopted genetic approach to perturb the membrane potential. Knocking out the genes (*nuoI*, *cydB*, and *aceE*) in the TCA and ETC pathways disrupted the most efficient energy production route, which would reduce the membrane potential ^23, 45^. Consistently, we found that the planktonic bacteria of these knockout strains grew better under TET stress than the WT strain did (Figures 6M and S7A). All perturbations above led to decreased membrane potential. To oppositely enhance membrane potential, we adopted the ion composition in the medium. In particular, when we increased the Na^+^: K^+^ ratio in the medium to cause cell membrane hyperpolarization in the biofilm, TET accumulation indeed increased significantly in the biofilm as predicted (Figures 6N and S7B-7F). We also explored the contribution of membrane hyperpolarization in biofilm to other drugs beyond TET. Similarly, CEF, CIP, and KAN accumulation and drug response increased significantly in the biofilm after the Na^+^: K^+^ ratio increased (Figures S7G-6L). Taken together, these results uncovered that the lower membrane potential functionally contributed to the slower accumulation of antibiotics in biofilms, causing less cellular damage in this critical time window, which enabled further adaptation including the induction of lipid metabolism, thickened cell membrane and in turn led to poorer drug penetration in biofilms. A model summarizing the molecular mechanism of biofilm antibiotic tolerance depending on such dynamic response was shown in Figure 7.

**Figure 7:**
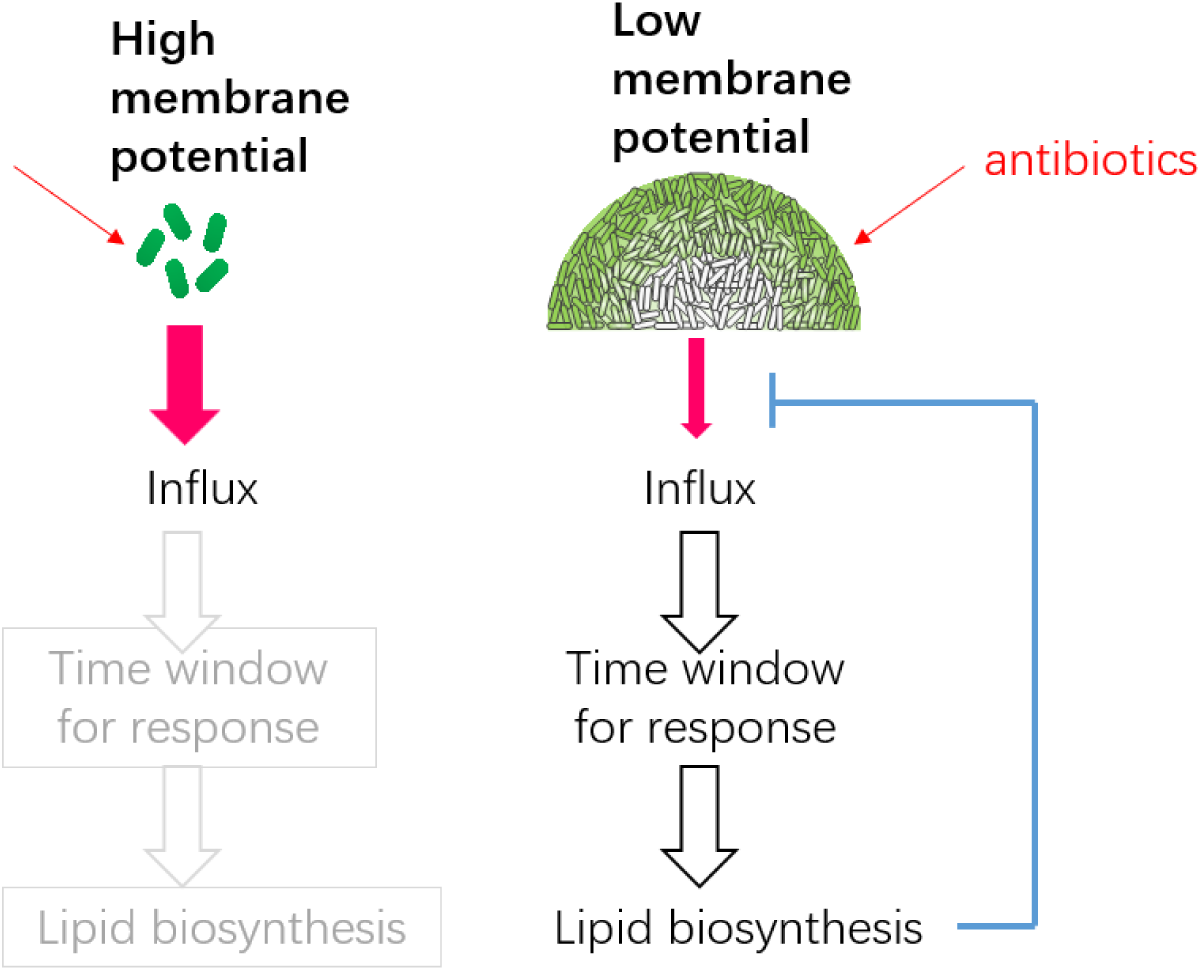
Antibiotic tolerance model for biofilm community.

## DISCUSSION

When compared to planktonic bacteria, biofilm communities exhibit spatial heterogeneity in terms of growth and metabolism, which may cause differences in the antibiotic enrichment and response to antibiotics. To date, the assumptions of biofilm tolerance to antibiotics is limited to the static aspects of community properties such as dormant subpopulations. Here, using advanced technologies including quantitative microfluidics, time-lapse microscopy and biofilm-based spatial multi-omics, we dissected the spatiotemporal drug response in biofilm. We found a common binary distribution for various antibiotics in biofilms, which was determined by glucose distribution. Antibiotics penetration would be accompanied by glucose leakage into the interior of the biofilm. Surprisingly, the biofilm periphery exhibited constant glucose consumption and antibiotics patterns after various antibiotics treatment, while planktonic bacteria did not. Thus, there was a common and biofilm-specific protection mechanism towards biofilm interior region adopted by different species to a wide range of antibiotics.

Spatial multi-omics revealed that activation of lipid metabolism was responsible for continuous glucose consumption, which was channeled to thicken the cell membrane and subsequently weakened antibiotic penetration, leading to a low drug concentration in the biofilm interior. We further revealed that the biofilm-specific activation of lipid metabolism depended on the inherently depolarized cellular membrane controlled by remodeled energy metabolism, biofilm limited drug penetration at the first stage, giving rise to a time window enabling constant glucose consumption to support further physiological adaptation. Our work also showed that changing the ion balance in the medium can promote the hyperpolarization of the biofilm, thus promoting the uptake of many antibiotics by the biofilm (Figures 6N and S7), which is a potential pharmaceutical strategy to reduce antibiotic tolerance of biofilm. The genetic program underlying the lower energy metabolism and membrane potential in the biofilm compared to planktonic bacteria (Figures 6I and 6J) is an important topic that needs exploration in future studies.

This study reveals that the unique activation of lipid metabolism is a common tolerance mechanism for a wide range of antibiotics in biofilms, and blocking the activation of lipid metabolism could effectively inhibit tolerance. Therefore, we assumed that the lipid metabolism pathway might serve as an effective drug target, especially for biofilms with inherently high drug tolerance. Interestingly, several chemicals have been found to inhibit key enzymes in this pathway ^46–48^, which would be promising to be explored in the future to eradicate biofilm-associated infections. Finally, the spatial proteomics in this study also uncovered that nucleotide metabolism was similarly activated in the biofilm under most antibiotic treatments (Figure 3B), which may also contribute to the antibiotic tolerance in a biofilm-unique manner. This is also of particular interest worthy further investigation.

## STAR★METHODS

### KEY RESOURCES TABLE

| Strain | Description |
| --- | --- |
| *E. coli* | BW25113 |
| *E. coli* with plasmid, pDL30 *PptsG-yfp* | *PptsG* was activated by glucose limited |
| *E. coli* with plasmid, pSB1C3 *PompF-mCherry* | *PompF* was activated by $\sigma_s$ and $\sigma_{70}$ |
| *E. coli* with plasmid, pSB1C3 *PtetR-mCherry* | *PtetR* was induced by aTc |
| *E. coli* with plasmid, pSB1C3 *Psyn213-mCherry* | A synthetic promoter being transcribed both by $\sigma_s$ and $\sigma_{70}$ <sup>49</sup> |
| *E. coli* with plasmid, pSB1C3 *Pcfa-mCherry* | *cfa* transcription was represented by mCherry fluorescence intensity <sup>49</sup> |
| *E. coli* with plasmid, $\Delta\psi$ sensor ViBac2 | Membrane potential $\Delta\psi$ distribution <sup>44</sup> |
| *E. coli* with knock out of *fabH* | Deficient in fatty acid biosynthesis initiation, from the Keio collection<sup>50</sup> |
| *E. coli* with knock out of *nuoI* | Deficient in NADH:quinone redox reaction, from the Keio collection<sup>50</sup> |
| *E. coli* with knock out of *cydB* | Deficient in NADH:quinone redox reaction, from the Keio collection<sup>50</sup> |
| *E. coli* with knock out of *aceE* | Deficient in pyruvate dehydrogenation, from the Keio collection<sup>50</sup> |
| *Klebsiella pneumoniae* | ATCC BAA-1144 |
| *Salmonella typhimurium* | ATCC 14028 |
| *Klebsiella pneumoniae* with knock out of *fabF* | Deficient in fatty acid biosynthesis initiation |
| *Salmonella typhimurium* with knock out of *fabH* | Deficient in fatty acid biosynthesis initiation |

| Reagents | Source |
| --- | --- |
| Tetracycline hydrochloride | Solarbio Life Sciences |
| Chloramphenicol | Solarbio Life Sciences |
| Ciprofloxacin hydrochloride | Solarbio Life Sciences |
| Ceftazidime | Sigma-Aldrich |
| Kanamycin sulfate | Sigma-Aldrich |
| Trimethoprim | Sigma-Aldrich |
| Rifampicin | Targetmol |
| Streptomycin sulfate | Targetmol |
| Anhydrotetracycline | Takara |
| CCCP | Sigma-Aldrich |
| FM4-64 (N-(3-triethylammoniumpropyl)-4-(p-diethylaminophenyl)hexatrienyl)-pyridinium 2Br) | Sigma-Aldrich |
| DiBAC4(5) | Life Technologies |
| 2-NBDG (2-[N-(7-nitrobenz-2-oxa-1,3-diazol-4-yl)amino]-2-deoxy-D-glucose) | Targetmol |
| DCFH-DA | Targetmol |
| [U-<sup>13</sup>C<sub>6</sub>]-D-glucose | Cambridge Isotope Laboratories |

| Antibiotics | Minimal inhibitory concentration of<br>*E. coli* Bw25113 in M63B1 (μg/ml) |
| --- | --- |
| Tetracycline hydrochloride | 2.0 |
| Chloramphenicol | 7.0 |
| Ciprofloxacin hydrochloride | 0.03 |
| Ceftazidime | 1.0 |
| Kanamycin sulfate | 5.0 |
| Trimethoprim | 5.0 |
| Rifampicin | 10.0 |
| Streptomycin sulfate | 5.0 |

### RESOURCE AVAILABILITY

#### Lead Contact

Further information and requests for resources and reagents should be directed to and will be fulfilled by the Lead Contact, Yuzhen Zhang.

#### Materials Availability

All bacterial strains are available upon request.

#### Data and Code Availability

All code used for data processing and analysis is available upon request. Any raw data not included in the repository, such as files in proprietary formats or large imaging files, are available upon request.

### EXPERIMENTAL MODEL AND SUBJECT DETAILS

On the day before the experiment, bacteria from −80℃ glycerol stock were streaked on LB agar plates and incubated at 37℃ overnight. On the next day, a single colony was picked from the plate, inoculated to 5 ml of LB broth in a 50 ml conical tube, and incubated at 37℃ in a shaker. After 12 h of incubation, the culture was centrifuged at a relative centrifugal force of 7000 g for 3 min. Then the pellet was re-suspended in 0.5 ml biofilm medium and used for loading into microfluidics. The biofilm and planktonic culture medium for *E. coli*, *S. typhimurium,* and *K. Pneumoniae* was standard M63B1 (100 mM KH_2_PO_4_, 15 mM (NH_4_)_2_SO_4_, 0.8 mM MgSO_4_, 3 μM vitamin B1, 22 mM glucose, adjusted to pH 7.4 with KOH) ^51^.

### METHOD DETAILS

#### Bacteria and cultivation system

##### Strains and plasmids

*Escherichia coli* strains: Promoters of *E. coli* strains (P*ptsG* and P*ompF*) were amplified from wild-type *E. coli* BW25113 genomic DNA; The P*tetR* sequence was synthesized by Genewiz (China). The promoter sequences were amplified by PCR and were then introduced to a plasmid containing fluorescent protein coding sequence to generate promoter activity reporters. Plasmids were constructed using the Gibson assembly method and transformed into DH5α competent cells (weidi, China). DNA purification and plasmid isolation were performed using Omega Bio-Tek (USA) reagents. PCR reactions were carried out using Q5 High-Fidelity DNA polymerase (NEB, UK). *Salmonella typhimurium* knockout strains used in this study are derivatives of strain 14028 (ATCC). Using Red/ET Recombination system, we constructed and verified *S. typhimurium* Δ*fabH*. The plasmids were pKD20, a Red helper plasmid containing a temperature sensitive origin of replication and ampicillin resistant marker which expresses recombinase enzyme in the presence of L-arabinose ^52^ and pKD4, a template plasmid which carries kanamycin gene flanked by FRT (FLP recognition target) sites ^53^. *Klebsiella pneumoniae* knockout strains used in this study are derivatives of strain BAA-1144 (ATCC). Using CRISPR-Cas9-mediated genome-editing method, we constructed and verified *K. pneumoniae* Δ*fabF*. A two-plasmid system, pCasKP-pSGKP, were used for precise and iterative genome editing. pCasKP-apr contains the Cas9 gene with a constitutive *rpsL* promoter and the temperature-sensitive replicon repA101(Ts) (repA101ts). pSGKP-km contains the sgRNA with the synthetic J23119 promoter and the *sacB* gene for plasmid curing ^54^.

##### Reagents

Tetracycline hydrochloride, chloramphenicol, and ciprofloxacin hydrochloride were purchased from Solarbio Life Sciences. Ceftazidime, kanamycin sulfate, trimethoprim, CCCP (carbonyl cyanide m-chlorophenylhydrazone) and FM4-64 (N-(3-triethylammoniumpropyl)-4-(p-diethylaminophenylhexatrienyl)-pyridinium 2Br) were purchased from Sigma-Aldrich. Anhydrotetracycline was purchased from Takara. DiBAC4(5) (Bis-(1,3-dibutylbarbituric acid) pentamethine oxonol) was purchased from Life Technologies. Rifampicin, 2-NBDG (2-[N-(7-nitrobenz-2-oxa-1,3-diazol-4-yl) amino]-2-deoxy-D-glucose) and DCFH-DA (dichloro-dihydro-fluorescein diacetate) were purchased from Targetmol. [U-^13^C_6_]-D-glucose was purchased from Cambridge Isotope Laboratories, Inc.

##### Microfluidic chip fabrication

The microfluidic chip was designed and manufactured in-house ^25^. The master mold of the microfluidic chip and spatial multi-omics chip was fabricated using the Maskless Mold Fabrication Method from BlackHole Lab (France) and performing lithography on silicon slides using a photoresist from Microchemicals (AZ4562). The microfluidic chips were made with polydimethylsiloxane (PDMS) and glass slides, while spatial multi-omics chips were only made with PDMS. The PDMS part was made by pouring a 10:1 (v/v) mixture of Sylgard 184 elastomer and curing agent (Dow Corning, USA) on the master mold, and curing the mixture for 2 hours in an oven at 80℃. Then, the cured PDMS with chamber pattern was carefully peeled off from the mold and punched with holes at the inlet and outlet ports. Then we pasted a (0.5-1 mm for microfluidic chip; 4-8 mm for spatial multi-omics chip) strip of 3M Scotch Magic tape to a designated location of the PDMS (on the barrier between the growth chamber and the loading channel, and perpendicular to the barrier), and treated the PDMS with plasma for 1-2 min (SoftLithoBox, BlackHole Lab, France). After the plasma treatment, the tape was quickly removed. The PDMS was immediately bounded with a glass slide (first-generation microfluidic chip) ^25^ or with a PDMS layer with a thickness of 1 mm (Figure 3A, spatial multi-omics chip). Finally, we plugged the inlet and outlet ports with syringe needles (20G, 0.91 mm OD x 0.61 mm ID) and PTFE Tubing (1/16” OD x 1/32 ID).

##### Cultivation of biofilm using microfluidic system

The cultivation of biofilm using microfluidics system followed our previous work ^25^. After the seed culture steps described before in experimental model, the culture was centrifuged at a relative centrifugal force of 7000 g for 3 min, and then the pellet was re-suspended in 0.5 ml PBS buffer, which served as the loading culture for microfluidics system. Elveflow OB1 Mk3 Pressure Controller (ELVESYS, France) was used to drive medium flow. At first, 5% BSA (bovine serum albumin) solution was used to treat the surface of microfluidics chamber for 30 min, so that the adhesion of individual bacteria onto the surface during loading would be inhibited; then switched back to PBS buffer. Subsequently, we loaded the abovementioned bacteria cells resuspended in PBS buffer, until ∼1000 bacteria cells were trapped at the seeding zone. During the rest of the experiment, the growth chamber was fed with biofilm medium under a constant pressure of 3 psi, and the temperature was kept at 37℃. The basic medium for biofilms was M63B1 ^51^. 10 μm 2-NBDG (which is imported and enriched in bacteria during glucose depletion ^23, 25^) or 5 μm DCFH-DA was added in the medium on demand. In the nutrient removing experiments, the biofilm medium was standard M63B1 without glucose. In the experiment of switching medium at different Na^+^/K^+^ ratios, there were 180 mM potassium ions in standard M63B1 and we adjusted Na^+^/K^+^ ratio by NaH_2_PO_4_ and NaOH under isosmotic conditions. In the experiment of biofilms response to antibiotics, the medium was switched to which contained 1-6 μg/ml tetracycline (TET), 10 μg/ml kanamycin (KAN), 14 μg/ml chloramphenicol (CM), 10 μg/ml streptomycin (Strep), 4 μg/ml ceftazidime (CEF), 0.06 μg/ml ciprofloxacin (CIP), 3 μg/ml trimethoprim (TMP), 20 μg/ml rifampicin (RIF), 200 or 400 μg/ml berberine (Berb), and 20 μg/ml chromomycin A3 (Chromo A3).

#### Microscopy

##### Time-lapse microscopy

The biofilms were observed with phase contrast and fluorescence microscopy. The microscope was Olympus IX83 (Japan) with Andor’s Zyla 4.2 sCMOS camera (UK). A 10X objective lens was used in most of the experiments to image the entire biofilm. Images were taken every 30 min, depending on the question. ***Confocal microscopy.*** Confocal imaging was performed using an inverted Zeiss LSM 710 three-channel laser scanning microscope (Carl-Zeiss) ^55^. The confocal microscopy samples were prepared by cultivation in our microfluidic system as described before. Then, biofilms were incubated in M63B1 containing 0.5 μg/ml FM4-64 fluorescent dye and 5 μg/ml TET for 20 h or 70 h. Finally, FM4-64 fluorescence was excited by a wavelength of 600 to 620 nm and detected at 650 to 683 nm ^31^; TET autofluorescence was excited at 405 to 420 nm and detected at 520 to 550 nm.

##### Transmission electron microscopy

TEM imaging was performed using a transmission electron microscopy (Tecnai Spirit 120kV). The TEM samples were prepared by cultivation in multi-omics microfluidic chip as described before. Then, biofilms were incubated in M63B1 containing 5 μg/ml TET for 20 hours or 60 hours and harvested in PBS. After washing twice in PBS, cells were collected on 3-mm copper (mesh) grids, fixed in 1% uranyl acetate and washed 3 times in ddH_2_O. Finally, the sample grids were dried in air for 3 minutes and the change of ultrastructure of bacteria membrane before and after TET treatment was observed under transmission electron microscopy. For each super-resolution image, the zoom factor of bacteria morphological characteristics images is 6800 x, zoom factor of bacteria membrane images is 30000 x. The final image was reconstructed using NIS-Elements software (Nikon) ^56^ (Figures 4D and S4B). The inner membrane was not always observed; therefore, it was not analyzed in this work.

#### Proteomic analysis

##### Proteome sample preparation

For planktonic cell samples, *E. coli* cells were grown in standard M63B1 medium at 37℃ under shaking at 220 rpm in 96-well plates and harvested in mid-exponential phase (OD_600_ = 0.25 ± 0.05). After reaching mid-exponential phase, *E. coli* planktonic cells were treated with antibiotics respectively (5 µg/ml TET, 10 µg/ml KAN, 3 µg/ml TMP, 14 µg/ml CM, 0.06 µg/ml CIP, 4 µg /ml CEF, 20 µg /ml RIF) for 20 h and harvested as planktonic cells antibiotics samples. For biofilm samples, *E. coli* cells were grown in standard M63B1 medium or treated with antibiotics for 20 h at 37℃ in our spatial multi-omics chip. The biofilm cultured in this microfluidic chip retained a semi-two-dimensional structure, and the upper and lower layers of the chip were made as thin polydimethylsiloxane (PDMS) layers with a thickness of 1 mm; thus, the chip appeared similar to a sandwich with two sliced pieces of bread. 30 min before the biofilm segmentation, we switched the medium including 10 μm 2-NBDG. Then we quickly separated the peripheral and interior regions of a biofilm along the 2-NBDG band under a stereofluoroscope (Figure 3A). Along with the upper and lower PDMS layers, we placed the two parts on ice. Then, we collected planktonic cell, biofilm interior and periphery samples by centrifugation at 13,000g at 4°C, washed twice with 200 μl ice-cold PBS buffer, harvested by centrifugation at 13,000g. Cells were resuspended in 100 µl lysis buffer (1% sodium deoxycholate, pH = 8.0 20 mM Tris - HCl) and heated to 90℃ for 10 min. The cells were disrupted by strong vortex followed by indirect sonication (100% amplitude, 45 s on 30 s off 20 cycles) in a bioruptorpico-2020. *E. coli* lysates were centrifuged at 16,000 x g for 15 min at 4℃ to remove cellular debris, the supernatants were collected and transferred to a fresh 1.5 ml tube and the protein concentration was determined by bicinchoninic acid assay (BCA kit, Thermo Fisher Scientific). The protein extracts were frozen in liquid nitrogen and stored at −80℃ until use. Then, the samples were diluted with lysis buffer to a final protein concentration of 0.5 mg/ml for each sample. Proteins obtained from the different samples were reduced with 5 mM TCEP for 60 min at 37°C and alkylated with 10 mM iodoacetamide for 30 min in the dark at 25°C. After quenching the reaction with light exposure for 20 min at 25°C, the samples were further digested by incubation with sequencing-grade modified trypsin (1/50, w/w; Promega, Madison, Wisconsin) overnight at 37°C. Protease digestions were quenched by lowering the reaction pH by adding wash buffer 1 (100% 2-isopropanol, 1% trifluoroacetic acid) to 4 times their original volume. The peptide mixtures were loaded onto SDB-Rps cartridges (Empore) and desalted. The desalted procedure was carried out with wash buffer 1 for one time and then wash buffer 2 (5% ACN, 0.1% trifluoroacetic acid) for one time to further remove the salt. The remaining peptides on cartridges were eluted with elution buffer (60% acetonitrile, 0.25% NH_3_ · H_2_O)^57^. After elution from the cartridges, peptides were dried in a vacuum centrifuge, resolubilized in 0.1% trifluoroacetic acid, and analyzed by mass spectrometry.

##### DDA Peptide pre-fractionation and spectral library generation

LC-MS/MS data dependent acquisition (DDA) peptide were fractioned by using HPLC (XBridgeTM BEH300 C18 column (Waters, MA)) to reduce sample complexity. For HPLC fractionation, buffer A was 100% H_2_O (pH = 10, adjusted by NH_3_·H_2_O) and buffer B was 98% acetonitrile (pH = 10, adjusted by NH_3_·H_2_O) using a HPLC gradient 72 min gradient (0-5 min, 5 to 8% of buffer B; 5-40 min, 8% to 18% of buffer B; 40-62 min, 18% to 32% of buffer B; 62-64 min, 32% to 45% of buffer B; 64-68 min, 95% of buffer B; 68-72 min, 95% to 5% of buffer B) at a flowrate of 1ml/min. The eluted peptides were ionized and detected by an Orbitrap Fusion mass spectrometry. The LC-MS/MS instrument used here was an UltiMate^TM^ 3000 RSLC nano system, directly interfaced with an Orbitrap Fusion LUMOS Tribrid mass spectrometer from Thermo Fisher Scientific. Peptides were loaded to a trap column (75 µm×20 mm, 3 µm C18,100 Å, 164535, Thermo Fisher Scientific) with a max pressure of 620 bar using mobile phase A (0.1% formic acid in H_2_O), then separated on an analytical column (50 cm length, 100 μm inner diameter, packed in house with ReproSil-Pur C18-AQ 1.9 μm resin from Dr. Maisch GmbH) with a gradient of 6-55% mobile phase B (80% acetonitrile and 0.08% formic acid) at a flow rate of 250 nL/min for 120 min. The MS data were acquired in data independent acquisition (DIA) mode and there was a single full-scan mass spectrum in the orbitrap (350–1650 m/z, resolution = 120, 000 at 200 m/z) with AGC target value of 2e6 and max injection time of 50 ms. Fragmentation was performed via a normalized collision energy of 35% with AGC of 5e5 and max injection time of 100 ms. Precursor peptides were isolated with 33 variable windows spanning from 350 to 1650 m/z at resolution=30,000. All the raw files were analyzed by Spectronaut 15.6 software. A spectral library (contains 2,923 E. coli proteins, 32,491 peptide sequences) was generated by DDA results with the search against Uniprot *E. coli* database (downloaded at 2021/07/01). Then the DIA results were analyzed using a targeted, library-based search with default settings. iRT (indexed retention time) reagent was used to normalize the retention time of peptides in all samples ^58^. Our DDA database consists of 2,923 *E. coli* proteins, resulting in a total of 32,491 protein sequences.

##### LC-MS/MS data acquisition

For LC-MS/MS analysis, samples were solubilized in solvent A (0.1% trifluoroacetic acid) at a concentration of 1 µg/µl. Subsequently, 1.2 µl 1*iRT reagent (Biognosys iRT Kit, Ki-3002-1) and 1.8 µl 0.01% trifluoroacetic acid were added into 5 µl samples and mixed thoroughly. Finally, 5 µl of the mix was injected per LC-MS/MS run. All samples of our data set were prepared in biological duplicates. Peptide separation was performed on a (75 µm × 45 cm) packed in-house with C18 resin (ReprosilAQ Pur, Dr. Maisch 1.9 µm) using a linear gradient from 95% solvent A (98% water, 2% acetonitrile, 0.1% trifluoroacetic acid) and 5% solvent B (98% acetonitrile, 2% water, 0.1% trifluoroacetic acid) to 30% solvent B over 120 min at a flow rate of 0.3 µl/min. For DIA sample measurements, 20 variable-width DIA isolation windows were recursively acquired. The DIA isolation setup included a 1 m/z overlap between windows, as described in previous research^59^. DIA-MS2 spectra were acquired at a resolution of 15,000 and an AGC target of 1 x 10^5^. To mimic DDA fragmentation, normalized collision energy was 32%, calculated based on the doubly charged center m/z of the DIA window.

##### Protein identification and label-free quantification

Targeted data extraction of DIA-MS acquisitions was performed with Spectronaut (Biognosys AG, version 13) with default settings, using spectral DDA libraries generated as described above. Briefly, the dynamic mass tolerance strategy was applied to calculate the ideal mass tolerances for data extraction and no correction factor was applied (correction factor = 1). The local (non-linear) regression method was used for iRT calibration using the iRT kit peptides. The relative quantitative data obtained were normalized and statistically analyzed using Spectronaut® Software ^26^, then we classified them according to Clusters of Orthologous Genes (COGs) ^27, 28^. The DIA dataset and COG classification results were shown in Supplementary table 1.

#### Metabolic and lipidomic analysis

##### 13C flux sample preparation and analysis

M63B1 medium with [U-^13^C_6_]-D-glucose was used in the ^13^C labeling experiments. First, we inoculated and cultivated the biofilm in standard M63B1. While we added antibiotics to treat biofilms in microfluidic chip, the medium was replaced by 100% [U-^13^C_6_]-D-glucose M63B1. Control samples were treated by changing to 100% [U-^13^C_6_]-D-glucose M63B1 and cultivated at the same time as the antibiotics treated group^60^. The cells were collected, washed, then resuspended into 80% (w/w) cold methanol, and disrupted in the same way as proteome samples. ^13^C labeling experiments samples, were analyzed on a Q Exactive™ Plus Orbitrap mass spectrometer (Thermo Fisher) with Ultimate 3000 Ultra High-Performance Liquid Chromatography (HPLC, Dionex Corporation of America). A Waters BEH Amide Column (2.1 × 100 mm, 1.7 μm) was used for HPLC separation. Buffer A was acetonitrile-0.1% formic acid and buffer B was water-10 mM NH_4_Ac-0.1% formic acid. HPLC gradient was 30 min (0-5 min, 0% of buffer B; 5-6 min, 0% to 25% of buffer B; 6-15 min, 25% of buffer B; 15-16 min, 25% to 50% of buffer B; 16-25 min, 50% of buffer B; 25-26 min, 50% to 0% of buffer B; 26-30 min, 0% buffer B) at a flowrate of 0.3 ml/min. Electrospray ionization was carried out by positive ion mode (spray voltage +3.2kV) and negative ion mode (spray voltage −3.0kV). For the MS1 full scan, ions with m/z ranging from 70 to 1000 were acquired by Orbitrap mass analyzer at a high resolution of 70,000. The automatic gain control (AGC) was set as 10^6^. The maximal ion injection time of full MS was 100 ms. MS2 acquisition was performed at 17,500 resolution and an AGC of 10^5^. Precursor ions were selected and fragmented with higher energy collision dissociation (HCD) with normalized collision energy of 30-40-50%. The maximal ion injection time of MS2 was 50 ms ^61^. Raw data were processed using Mzvault 2.2 database (Thermo Fisher) and Xcalibur Qual Browser (Thermo Fisher). For each scan mode, metabolite identities were confirmed using authentic reference standards from IROA Technologies Company. Metabolite molecular formulas matching to within 5 parts per million (ppm) were selected as candidate identifiers. Identification of unknown compounds using compound discovery 3.2, mzcloud, and mzVault databases ^62^.

##### Lipidomics sample preparation and analysis

Samples for lipidomics analyses were collected, washed, and harvested with the same processes as proteome samples we describe above. Total fatty acids were extracted from cell cultures by chloroform/methanol (2:1 vol/vol) as previously described ^61^. The cells were disrupted by strong vortexing followed by indirect sonication (100% amplitude, 45 s on 30 s off 20 cycles) in a bioruptorpico-2020. After 5 min of low-speed centrifugation, the upper phase was discarded, and the lower phase (containing the lipid extract) was collected. Re-extracted upper phase with 500 µL methanol, and 200 µL water and collected the organic phase (top layer) again and combined the two organic phases. All collected lipid extracts were brought to dryness using a gentle stream of nitrogen ^63^. The lipid extracts were resuspended by 50 μl chloroform/ isopropanol (2:8 vol/vol) The lipid fractions in sample were subsequently analyzed by separate liquid chromatography-mass spectrometry (LC-MS) methods for quantification. We combined Ultimate 3000 Ultra High-Performance Liquid Chromatography (Dionex Corporation of America) with Thermo Q Exactive Plus High-Resolution Mass Spectrometry (Thermo Fisher Scientific of the United States). Lipid samples were injected and separated on an ACQUITY UPLC BEH C18 Column (2.1 × 100 mm, 1.7 μm). For HPLC separation, buffer A was 60% acetonitrile-40%H_2_O-10 mM NH_4_Ac-0.1% formic acid and buffer B was 80% isopropanol-20% acetonitrile-10 mM NH_4_Ac-0.1% formic acid. HPLC gradient is a 60 min gradient (0-5 min, 15 to 30% of buffer B; 5-10 min, 30% to 50% of buffer B; 10-15 min, 50% to 54% of buffer B; 15-20 min, 54% to 70% of buffer B; 20-35 min, 70% to 99% of buffer B; 35-45 min, 99% B; 45-50 min, 99% B to 15% B; 50 - 60 min, 15% B) at a flowrate of 0.25 ml/min. Electrospray ionization were carried out by positive ion mode (spray voltage +3.2kV) and negative ion mode (spray voltage −3.0kV). For the MS1 full scan, ions with m/z ranging from 100 to 1500 were acquired by Thermo Q Exactive mass analyzer at a high resolution of 70,000. The automatic gain control (AGC) was set as 10^6^. The maximal ion injection time of full MS was 100 ms. MS2 acquisition was performed at 17,500 resolution and an AGC of 10^5^. Precursor ions were selected and fragmented with higher energy collision dissociation (HCD) with normalized collision energy of 30-40-50%. The maximal ion injection time of MS2 was 50 ms. Measure protein concentration in each sample as normalization of metabolites abundance by resuspending with 8 M urea buffer and then quantified using the BCA Protein Assay Kit described above. A dataset about lipidomics was shown in Supplementary table 2.

#### Glucose and antibiotics quantification

##### Intracellular antibiotics concentration measurement

Samples for antibiotics analyses were collected, washed, and harvested with the same processes as the proteome samples described above. Antibiotics were extracted from samples with 80% methanol. The antibiotics fractions (TET, CIP, CEF, and Berb) in the sample are subsequently analyzed by HPLC (Agilent 1290) methods for quantification. Antibiotics samples were injected and separated on a Bridge Shield RP18 Column (4.6 × 250 mm, 5 μm). A gradient of buffer A (0.1% formic acid) and buffer B (acetonitrile) was used at a 1 ml/min flow rate. HPLC gradient is a 25 min gradient (0-5 min, 0 to 10% of buffer B; 5-8 min, 10% to 20% of buffer B; 8-15 min, 20% to 80% of buffer B; 15-25 min, 80% to 10% of buffer B). 5 µl of each sample was injected for analysis. We combined HPLC (Agilent 1290) with Mass Spectrometry (Agilent 6460 Triple Quad LC/MS) for KAN quantification. Antibiotics samples were injected and separated on an XSelect HSS T3 Column (2.1 × 50 mm, 2.5 μm). A gradient of buffer A (0.1% formic acid) and buffer B (acetonitrile - methanol - 0.1% formic acid) was used at a 0.2 ml/min flow rate. The HPLC gradient is a 10 min gradient (0 - 7 min, 0% to 20% buffer B; 7 - 9 min, 20% to 100% buffer B; 9 - 10 min, 100% to 20% buffer B).

##### Extracellular glucose concentration measurement

Glucose (GO) Assay Kit (Sigma-Aldrich) was used to quantify the extracellular glucose concentration in inlet standard M63B1 medium, waste of biofilm samples from outlet tube, and supernatant of planktonic bacteria (before and after antibiotics treatment for 3 h, 6 h, 9 h, 12 h, and 15 h). After centrifugation (10 min, 13,000g at 4°C), the biofilm and planktonic bacteria samples were diluted with PBS by 1:100. The glucose concentration of dilution samples was quantified by glucose (GO) Assay Kit (Sigma-Aldrich) and measured by a microplate reader (TECAN, Swiss) ^64^.

##### Membrane hyperpolarization and damage measurement

DiBAC4(5), which is a negative lipophilic dye used to detect membrane potential and membrane damage, and exhibits enhanced fluorescence during membrane depolarization and membrane damage ^24, 41^. After the seed culture steps described before in experimental model, bacterial culture was pelleted at 7000 x g for 3 min and washed twice in PBS. Then, the pellet was re-suspended by 10 ml M63B1 medium and 200 μl of the culture was inoculated to 10 ml M63B1 medium containing 500 μM DiBAC4(5) and incubated in a dark 37℃ shaking incubator for 1 h. After incubation, we loaded the culture into the microfluidic chip, and DiBAC4(5) fluorescence was measured by microscopy at the RFP channel (590 nm excitation and 620 nm emission wavelengths) ^65^. A time-course acquisition was performed, with 5 μg/ml TET injected to the chamber to measure increases or decreases of DiBAC4(5) fluorescence. Besides, the biofilms were first cultured in M63B1 medium containing 500 nM DiBAC4(5), then the medium containing 500 nM DiBAC4(5) and 5 μg/ml TET was switched after the biofilm size reaching to 500 μm.

##### Planktonic bacteria growth curve and CFU experiments

The culturing of planktonic *coli* (*ΔnuoI, ΔaceE, ΔcydB,* and WT strains) were performed using the SPARK microplate reader (TECAN, Swiss). The bacteria were grown overnight in the LB medium in a 37°C shaker. On the next day, 4 μL of the culture was inoculated to 200 μL of fresh M63B1 and incubated in 96-well plates. When the cells grown to log phase (OD_600_ = 0.2∼0.3), 1 μg/ml TET was added in the medium (Figure S7A). Growth curves about optical density (600 nm) was recorded every 20 min. All 96-well experiments were performed with biological replicates from independent cultures. For colony forming unit (CFU) experiment, when the cells grown to log phase (OD_600_ = 0.2∼0.3), bacteria cultures were centrifuged, resuspended by different volume PBS, and adjusted to same final bacterial concentration (OD_600_ = 0.1). Then, bacteria cells were diluted 1:1000 by PBS and spread on M63B1 plate containing 1μg/ml TET. After 18 h cultivated in a 37°C incubator, bacteria numbers on plates were quantified by CFU (Figure 6M).

##### FACS experiment

FACS experiments was performed in V500-A channel on a FACS Aria SORP (BD Biosciences). A blank *E. coli* Bw25113 strain was used as a negative control in the primary cytometry setting. After growth to mid-exponential phase (OD_600_ = 0.3) in SPARK microplate reader, the cultures were treated under 5 μg/ml TET for different time durations (1 h, 2 h, 3 h, 4 h, and 5 h). Then, we chilled the samples with or without TET on ice immediately and diluted in 1 ml of prechilled PBS. A 500-fold dilution was applied to each sample, making them appropriate samples for FACS experiments.

##### Image analysis

ImageJ (National Institutes of Health, USA) and MATLAB (MathWorks, USA) were used for data analysis. For each snapshot of the biofilm from time-lapse microscopic images, we cut a line (1 pixel wide) along the direction of biofilm expansion as the region of interest (ROI) (Figure S1A), and put all ROIs into a kymograph according to time sequence. On the phase-contrast kymograph (Figure S1B), we set an appropriate threshold to find the edge of the biofilm. The information is merged with the fluorescent kymograph (Figure S1D) to obtain the terminal kymograph as shown in Figure S1F. The terminal kymograph (Figure S1F) contains almost all the spatiotemporal information of the biofilm (size, growth, and fluorescence changes before and after the perturbation). Based on this image (Figure S1F), we further extract the spatial profile of fluorescence intensity at a specific time, also extract the temporal profile of average fluorescence in a region with an equal distance (blue lines and green line) from the edge of the biofilm (Figure S1G and 1H).

## Acknowledgements

We thank Jintao Liu for technical supports for development of microfluidics for biofilm culturing and quantitative analysis of data, thank Fan Bai in Peking University for kindly providing bacterial strains (membrane potential sensor ViBac2), thank Shouxian Hu and Ziyan Wang for data analysis, thank the faculty of Protein Chemistry and Proteomics Platform at Tsinghua University for assistance with proteomics assays, thank Tsinghua University Center of Pharmaceutical Technology for assistance with metabolomics assays. Y.Z. was supported by the National Nature Science Foundation of China (21908129), Chinese Postdoctoral Science Foundation (2018M631481), and the Tsinghua-Peking Center for Life Sciences.

## Author contributions

Y.Z. designed the study, Y.Z. designed the microfluidic chip, Y.Z., Y.C. and Q.W. performed the experiments, Y.Z., Y.C. and T. W analyzed the data, Y.Z., Y.C. and T. W wrote the manuscript, all authors discussed the manuscript.

## Declaration of interests

The authors declare no conflict of interest.

## Supplementary Figures

**Figure S1:**
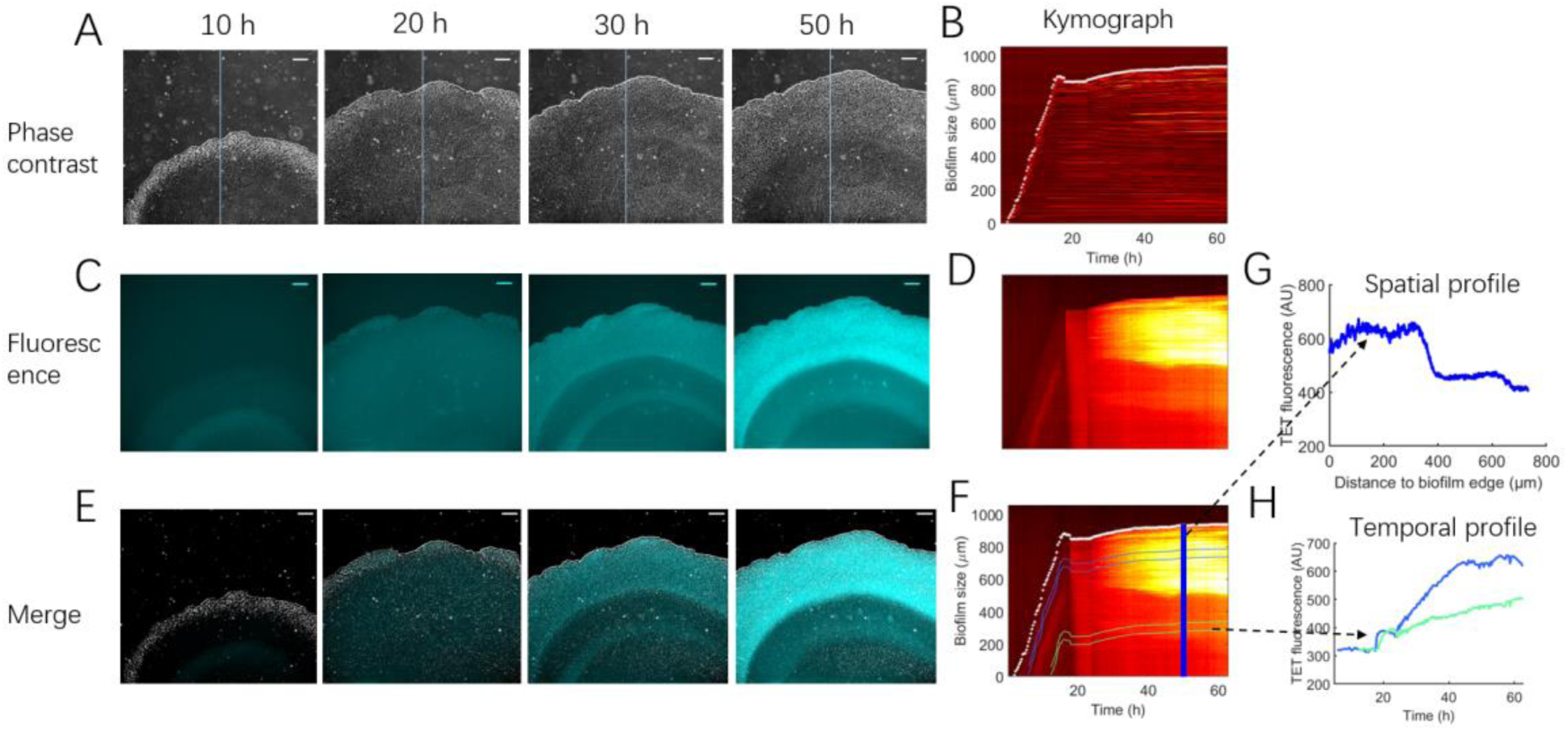
Image processing. (A, C, and E) Snapshots of phase-contrast, fluorescent, and merged images for *E. coli* biofilm. (B, D, and F) Phase-contrast, fluorescent, and merged kymograph for *E. coli* biofilm. (G) The spatial profile of fluorescence intensity at a specific time. (H) The temporal profile of average fluorescence in biofilm periphery (blue lines) and interior (green line) with an equal distance to the edge of the biofilm.

**Figure S2:**
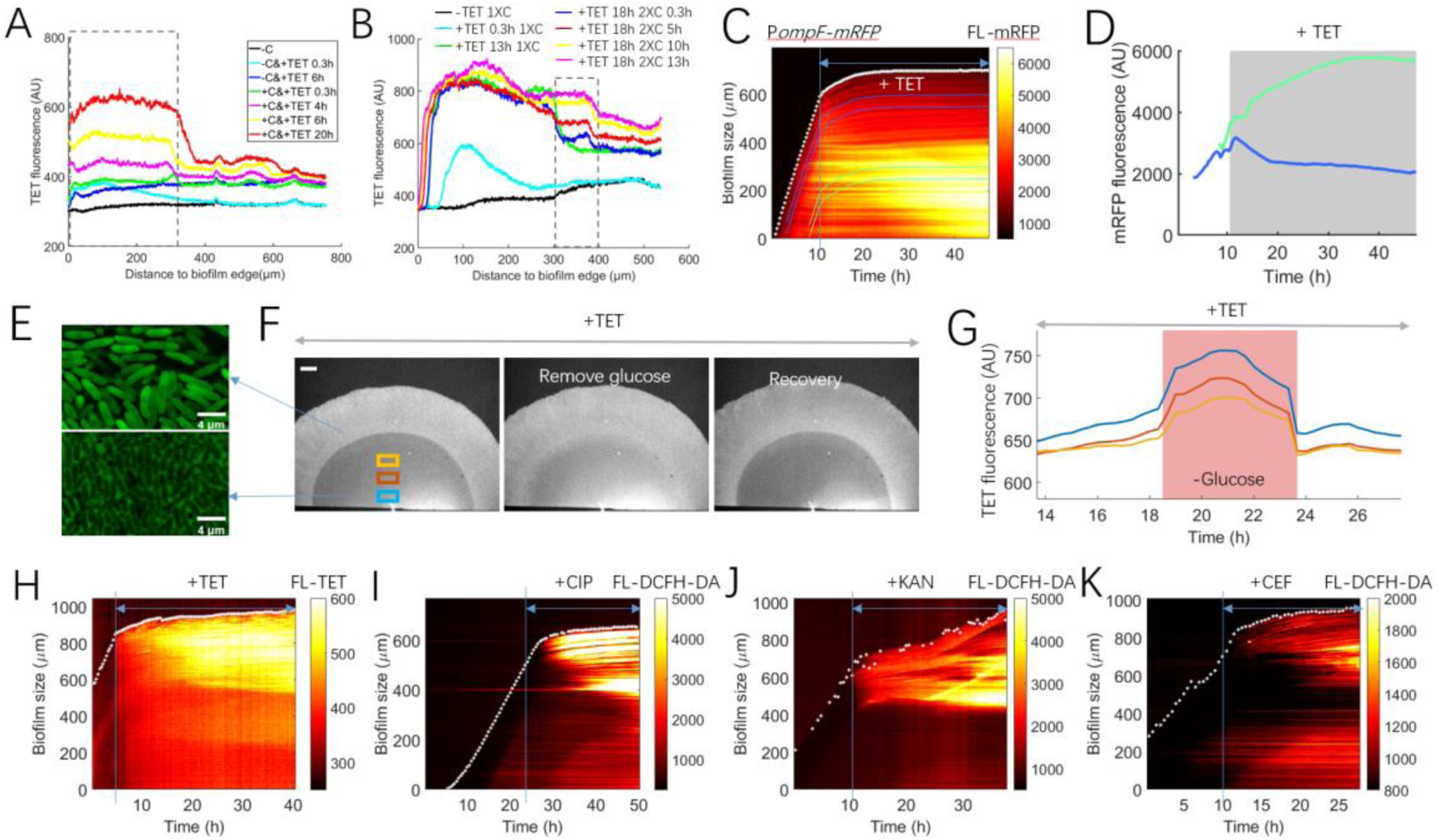
Antibiotics enrichment in *E. coli* biofilm. (A and B) The profile of tetracycline (TET) was changed when glucose concentration in the medium was perturbed. (C and D) mRFP translation in P*ompF*-mRFP biofilm before after addition of 5 μg/ml TET (periphery, blue line; interior, green line). (E) 5 μg/ml TET accumulation in the periphery (top) and the interior (bottom) of the biofilm. (F and G) The TET (5 μg/ml) fluorescence in the interior of the biofilm before and after removing glucose from the medium. (H-K) Kymograph of antibiotics accumulation in the biofilm. 5 μg/ml TET, 0.06 μg/ml ciprofloxacin (CIP), 10 μg/ml kanamycin (KAN), and 4 μg/ml ceftazidime (CEF) were added.

**Figure S3:**
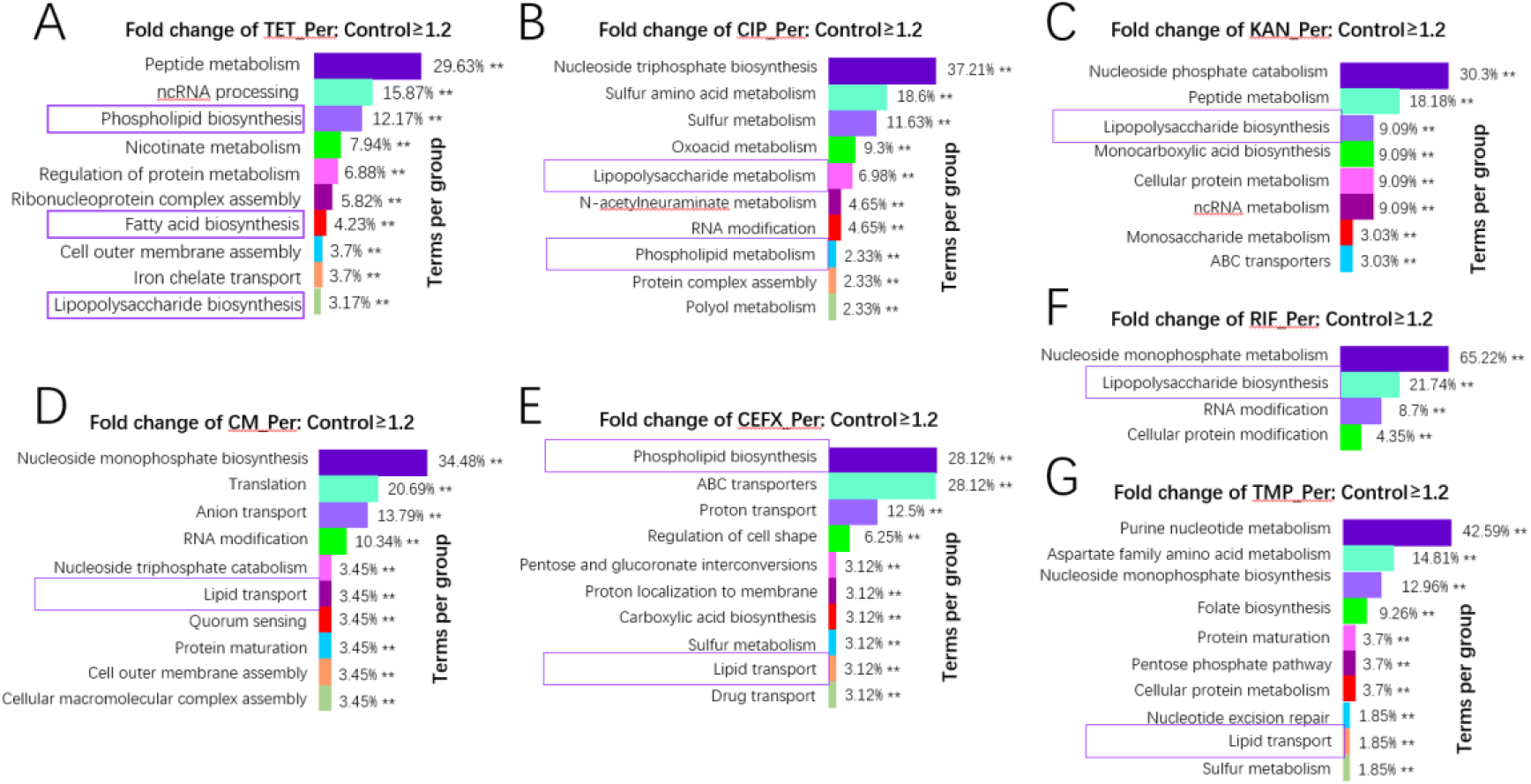
Pathway enrichment for proteins in biofilm periphery with fold-change of antibiotics treatment /no treatment ≥ 1.2. p ≤ 0.05.

**Figure S4:**
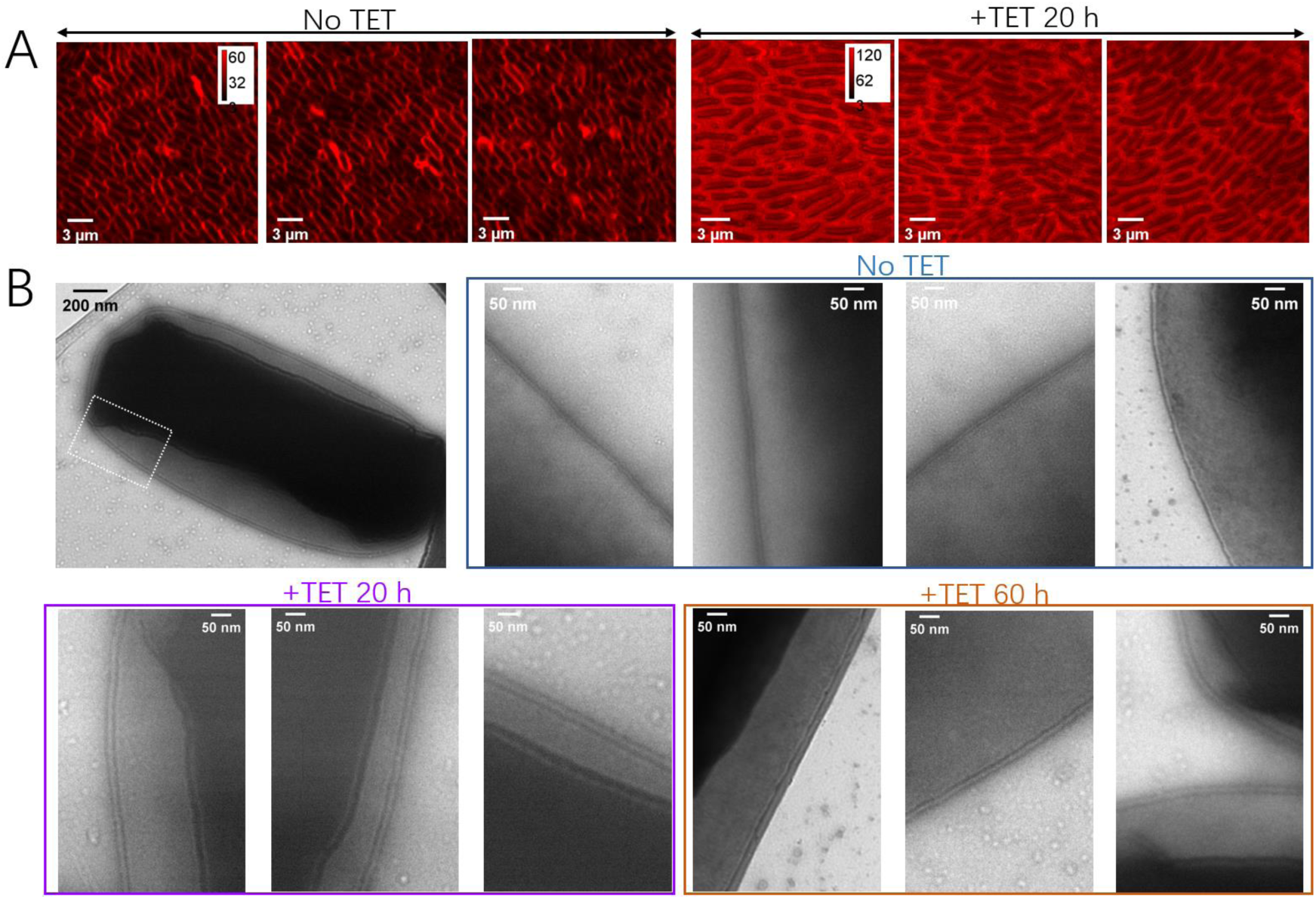
Cell envelope before and after tetracycline (TET) treatment. (A) Confocal images of the cells in the biofilm periphery with (20 h) or without 5 μg/ml TET. 0.5 μg/ml FM 4-64 was added. (B) Transmission electron microscopy images of the cells in the biofilm periphery with (20 h and 60 h) or without 5 μg/ml TET.

**Figure S5:**
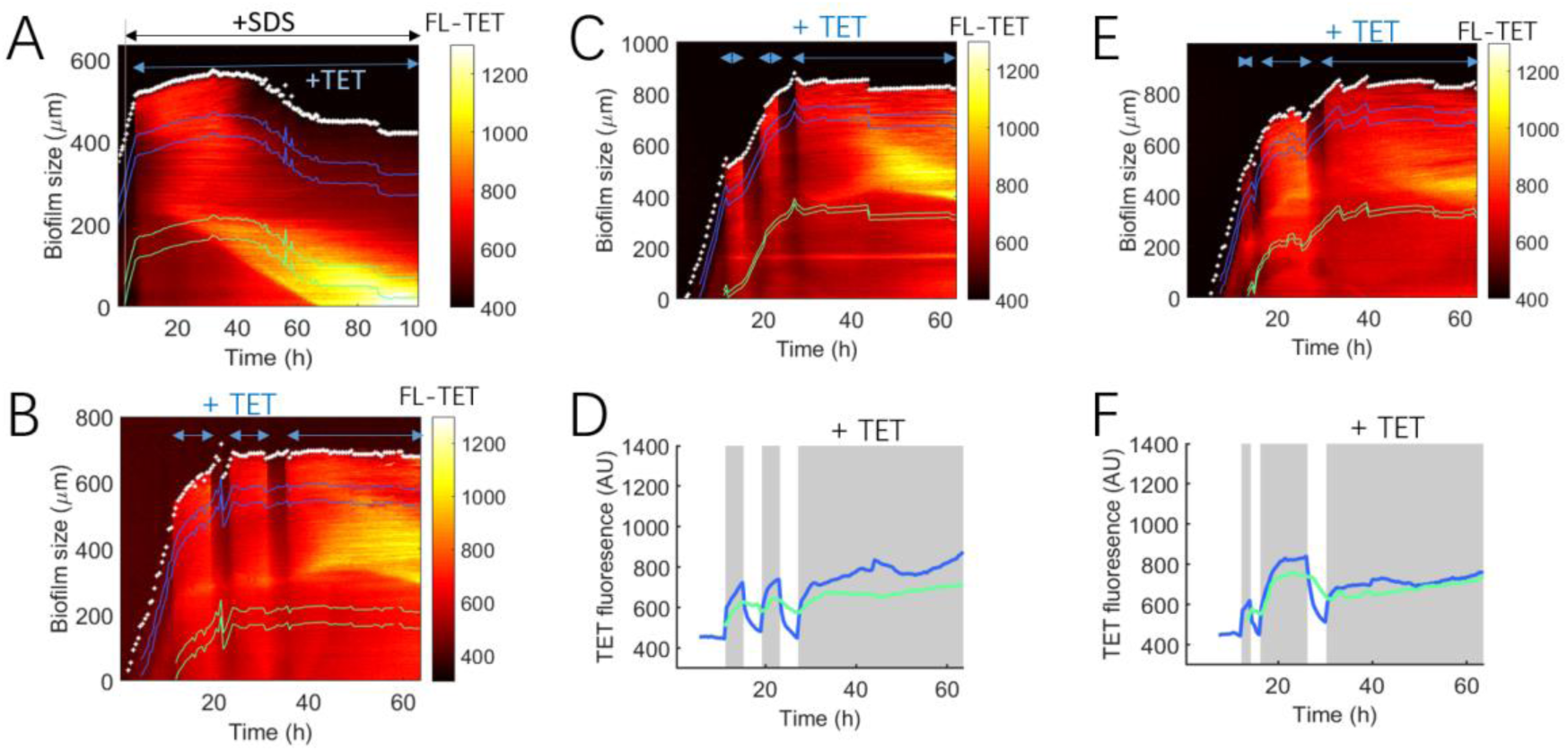
Tetracycline (TET) enrichment in *E. coli* biofilm. (A) Kymograph of 5 μg/ml TET pattern in WT *E. coli* biofilm when 0.03% sodium dodecyl sulfate (SDS) was added 2 h in advance. (B, C and E) Kymograph of 5 μg/ml TET pattern in WT *E. coli* biofilm when TET was added periodically. (D and F) TET fluorescence accumulation in biofilm (periphery, blue line; interior, green line) when 5 μg/ml TET was added periodically shown in Figure S5C and 5E, respectively.

**Figure S6:**
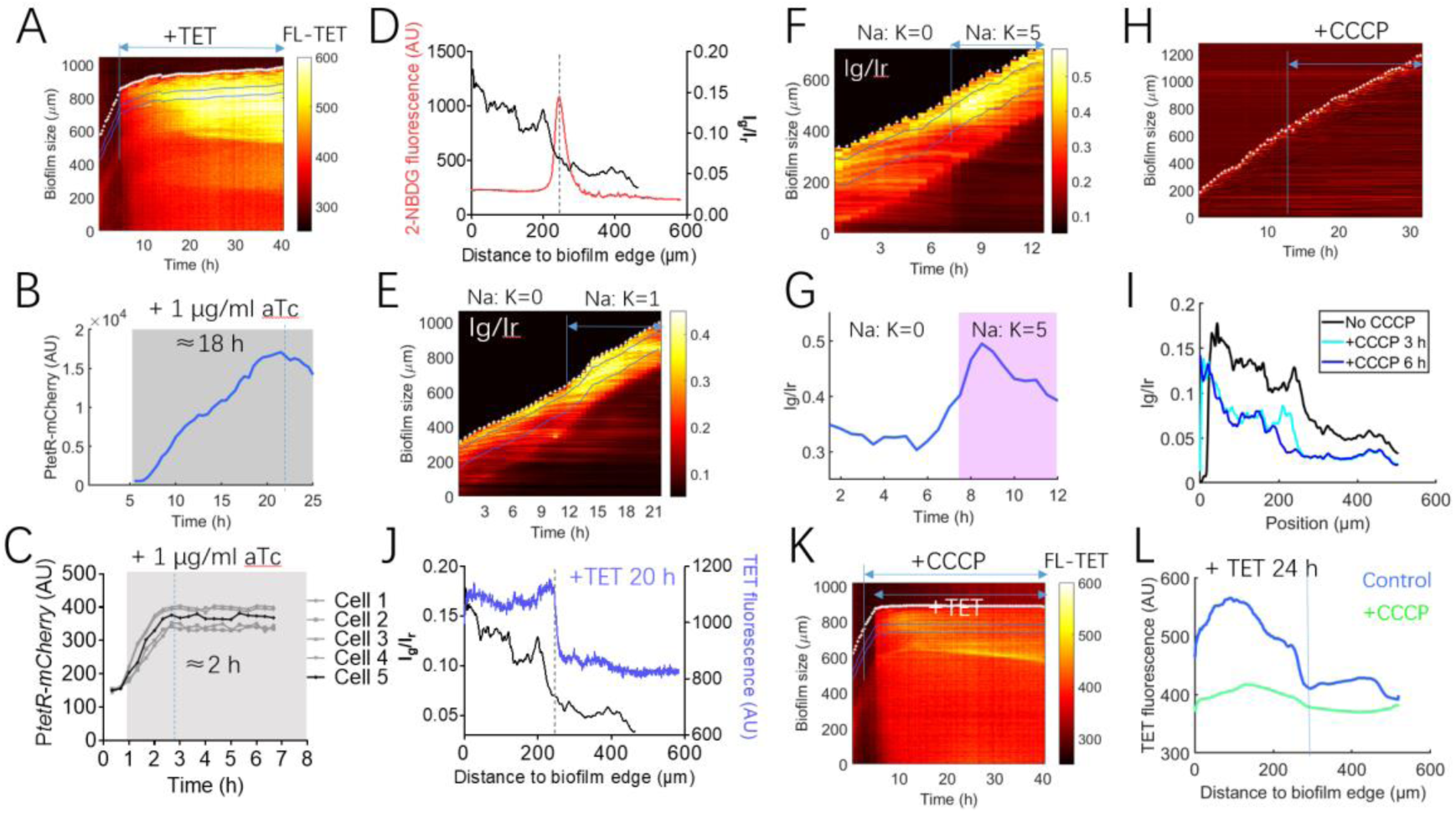
Tetracycline (TET) enrichment in bacteria and caused by membrane potential. (A) Kymograph of TET pattern in WT *E. coli* biofilm. 5 μg/ml TET was added. (B and C) mCherry fluorescence in P*tetR*-mcherry biofilm (B) and planktonic cells (C) after the addition of 1 μg/ml anhydrotetracycline (aTc). (D) The distribution pattern of membrane potential was highly consistent with that of 2-NBDG. (E and F) Kymograph of membrane potential in WT *E. coli* biofilm before and after changing the Na^+^: K^+^ ratio. (G) Membrane potential (Ig/Ir) of cells in the biofilm increased significantly after changing the Na^+^: K^+^ ratio from zero to 5 shown in (F). (H) Biofilm growth was not affected by 20 μM CCCP. (I) Spatial profile of membrane potential before and after addition of 20 μM carbonyl cyanide m-chlorophenyl hydrazone (CCCP). (J) The distribution pattern of membrane potential was highly consistent with that of TET. (K) Kymograph of TET pattern in WT *E. coli* biofilm when 20 μM CCCP added. (L) The spatial distribution of TET under 20 μM CCCP added or not.

**Figure S7:**
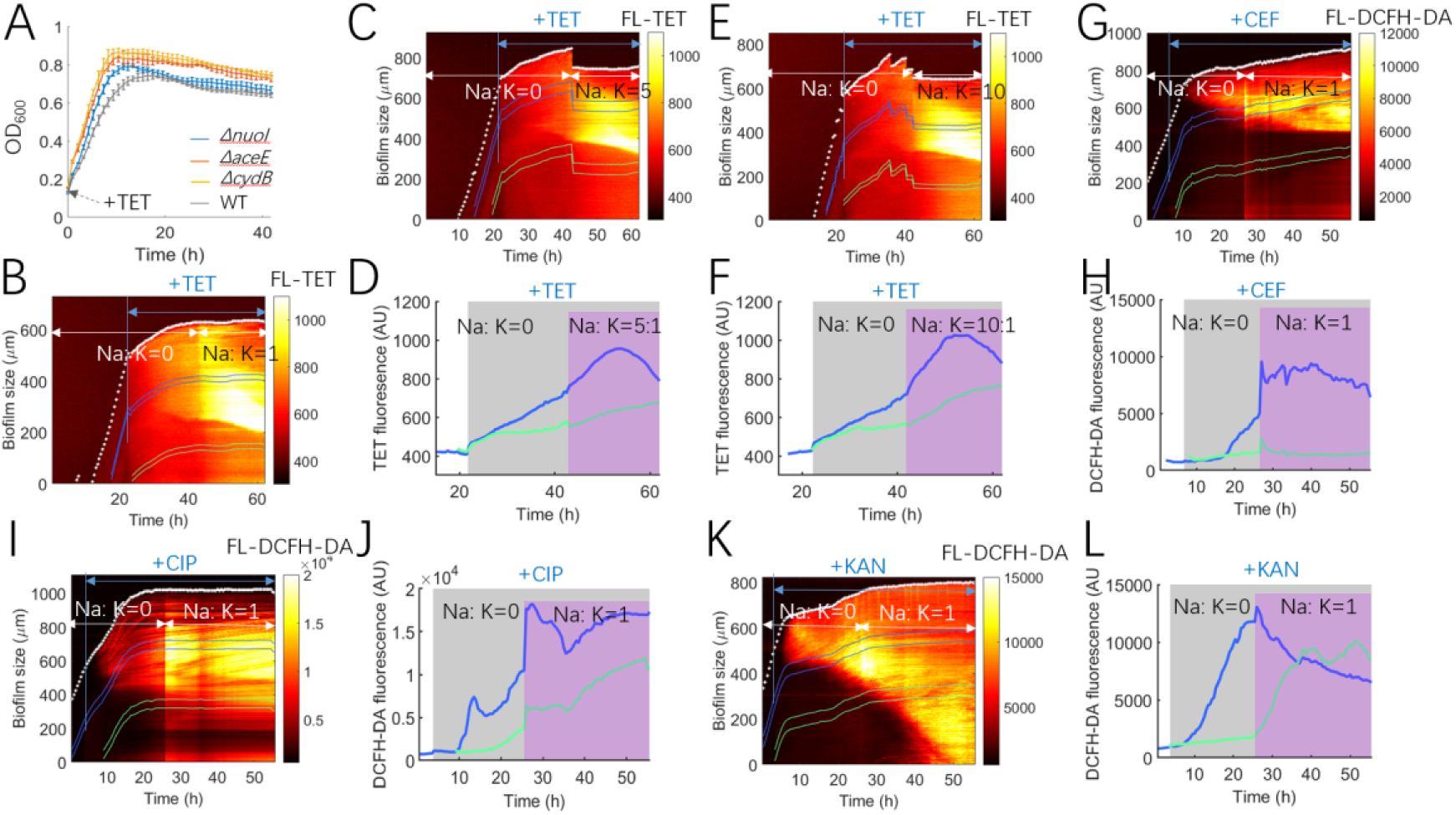
The change of membrane potential of biofilm resulted in the drug enrichment. (A) Growth curves of *ΔnuoI, ΔaceE, ΔcydB,* and WT strains under 1 μg/ml tetracycline (TET). Zero time represents when TET was added. (B, C, and E) Kymograph of 5 μg/ml TET in WT *E. coli* biofilm before and after changing the Na^+^: K^+^ ratio. (D and F) TET fluorescence increased significantly after changing the Na^+^: K^+^ ratio after adding TET shown in (C) and (F) respectively. (G, I, and K) Kymograph of various antibiotics (ceftazidime (CEF, 4 μg/ml), ciprofloxacin (CIP, 0.06 μg/ml), and kanamycin (KAN, 10 μg/ml)) in WT *E. coli* biofilm before and after changing the Na^+^: K^+^ ratio from zero to one. (H, J, and K) Accumulation of various antibiotics increased significantly in the biofilm after the Na^+^: K^+^ ratio increased shown in (G), (I), and (K), respectively.

## Notes

### Competing Interest Statement

The authors have declared no competing interest.

